# Genetically restricted human antibodies recognize a conserved β-hairpin in postfusion influenza hemagglutinin

**DOI:** 10.64898/2026.10.28.760416

**Authors:** Joel Finney, Juyeon Park, Martin Dodes Traian, Masayuki Kuraoka, Dalton Towers, M. Anthony Moody, George Georgiou, Garnett Kelsoe, Stephen C. Harrison

**Author notes:** Correspondence: Joel Finney,; Stephen C. Harrison.

## Abstract

A public class of broadly protective, human antibodies (Abs) binds a conserved β-hairpin exposed in the postfusion conformation of influenza virus hemagglutinin (HA). We here describe use of *in vitro* immunoassays and high-resolution cryogenic electron microscopy analysis of Ab:HA complexes to explain the apparent genetic restriction of these Abs and the wide range of influenza A and B subtypes they recognize. We show that recognition of the β-hairpin epitope requires 1) germline-encoded residues unique to certain alleles of the human immunoglobulin heavy chain gene *IGHV1-2*, 2) a short (<14-residue) HCDR3 to avoid steric clash with HA, and several Ab light chain residues that are germline-encoded by various *IGLV2-*family genes or *IGLV7-46*. Despite the genetic restrictions, the capacity to make β-hairpin Abs is widespread among humans. We recovered examples of these Abs from 14 genetically unrelated donors, and a previously reported analysis of the *IGHV* genotypes of ∼2,500 individuals sampled from major ethnic populations estimates that >75% of people have the necessary *IGHV1-2* alleles to make these Abs. We conclude that an immunogen that displays the β-hairpin epitope might reliably elicit broadly protective influenza antibodies in most humans.

## INTRODUCTION

Influenza A and B viruses (IAVs/IBVs) cause substantial morbidity and mortality in human populations (*1*), with accompanying economic burden (*2*). The phylogenetically and antigenically diverse IAVs are classified by differences in their surface glycoproteins: hemagglutinin (HA), which mediates viral attachment and membrane fusion with the host cell; and neuraminidase (NA), which enables viral egress from the cell (*3*). Various combinations of 16 subtypes of HA (H1-H16) and 9 subtypes of NA (N1-N9) represent all non-bat IAVs (*4*); the H17 and H18 subtypes have been found only in bats (*5*). IBVs comprise two lineages, Victoria and Yamagata. B/Victoria and H1N1 and H3N2 IAVs currently circulate among humans, while additional pre-pandemic strains circulate in animal reservoirs, occasionally causing severe disease in humans (*6, 7*).

Seasonal flu vaccines offer modest protection from disease, but rely on predictions about which viral strains will predominate during the coming flu season (*8, 9*). Antigenic mismatches between the predicted and circulating strains frequently occur, lowering vaccine effectiveness (*8, 10*). Population-level immune pressure drives antigenic “drift”, *i.e.*, selection of HA and NA mutations that promote viral escape from immune control (*11, 12*). Consequently, annually updated, licensed influenza vaccines typically protect only against seasonal strains and closely related subtypes (*9, 13*). In addition, the sporadic transmission of influenza viruses from animal reservoirs to humans can obviate pre-existing immunity, causing pandemics (*14*). Improved flu vaccines are clearly needed.

Rare Abs to conserved epitopes in HA protect against disease by diverse IAVs and IBVs (*15–18*). Among these Abs is a public class of human Abs to a conserved β-hairpin in the HA2 subunit; this β-hairpin is at least partly occluded in prefusion HA but fully exposed in postfusion HA (*19*). HA β-hairpin Abs recognize antigenically-drifted IAVs and IBVs that have circulated among humans for at least the last 68 years (*19*), as well as many pre-pandemic flu viruses (*19*). All five known β-hairpin Abs have heavy chains (HCs) encoded by *IGHV1-2* and light chains (LCs) derived from *IGLV2-11* or *IGLV2-23* (*19*). Although β-hairpin Abs are non-neutralizing, they recruit Fc effector activities to protect mice from lethal influenza challenge (*19*), as do Abs against other conserved, non-neutralizing HA epitopes (*18, 20–23*). A previous (*19*), ∼5Å-resolution cryogenic electron microscopy (cryo-EM) reconstruction of a representative β-hairpin Fab bound to a fragment of postfusion HA2 could not explain why β-hairpin Abs are apparently genetically restricted, nor how these Abs bind so many HA subtypes. Also unresolved is whether humans are widely capable of mounting humoral responses to the HA2 β-hairpin. Answers to these questions, provided here, indicate that vaccination with postfusion HA could reliably elicit HA2 β-hairpin Abs in humans and that such Abs could contribute to the broad, durable protection desired from next-generation influenza vaccines.

## RESULTS

### Prospective identification of HA2 β-hairpin monoclonal Abs (mAbs) from ten new donors

On the basis of *in vitro* HA-binding assays and/or V(D)J sequence similarities to known β-hairpin mAbs (*19*), we identified 34 potential HA2 β-hairpin mAbs from eleven donors not previously known to have humoral responses against the β-hairpin (BAA001, O100, B5, C1, C4, P04, P05, P11, 310, 014-10, 019-10) and from one donor already known to make β-hairpin Abs (donor 350). Some of these mAbs had been identified by us (*24*)(Towers *et al.*, in preparation) or others (*25, 26*) as HA-reactive Abs with unknown epitopes, while the remaining mAbs had not been checked previously for HA-reactivity (*27–29*). We tested whether recombinant IgG forms of these mAbs could bind recombinant HA2 in its postfusion conformation (EHA2) (*30*) in ELISA. Twenty-one IgGs did not bind EHA2s (**Table S1**). The remaining 13 IgGs competed with the archetypal β-hairpin mAb, S1V2-72, for binding to EHA2, but they did not compete with control mAb CR9114 (**Fig. 1A**). The 13 newly identified β-hairpin IgGs and six previously reported β-hairpin mAbs (S1V2-72, K06.18, 334-100, 350-310-D7B, 350-315, 350-376) (*19*) each bound EHA2s representing the H1 and H3 subtypes of seasonal IAVs, several other pre-pandemic IAVs that circulate in bird populations, and IBV (**Fig. 1B, Fig. S1A**). Some of these mAbs also bound full-length, soluble ectodomain (FLSE) versions of H17 and/or H18, which represent IAVs that circulate among bats. (Although FLSEs assume the prefusion conformation in solution, adsorption to ELISA plates seems to expose epitopes that are normally occluded in prefusion HA, including the conformational epitope bound by LAH31, which otherwise is revealed only in postfusion HA2 (*23*)). None of the mAbs bound EHA2s in which the tip of the β-hairpin loop had been mutated from N135_HA2_ to R135_HA2_ (N135R)(**Fig.1B, Fig. S1B**). In contrast, control mAbs LAH31 (*23*) or S5V6-P7F12 (manuscript in preparation), which do not bind the β-hairpin, bound wildtype (WT) and N135R EHA2s equally well (**Fig. 1B, Fig. S1A-S1B**). Control mAb CR9114 did not bind EHA2s, but did bind FLSE HAs.

**FIGURE 1.**
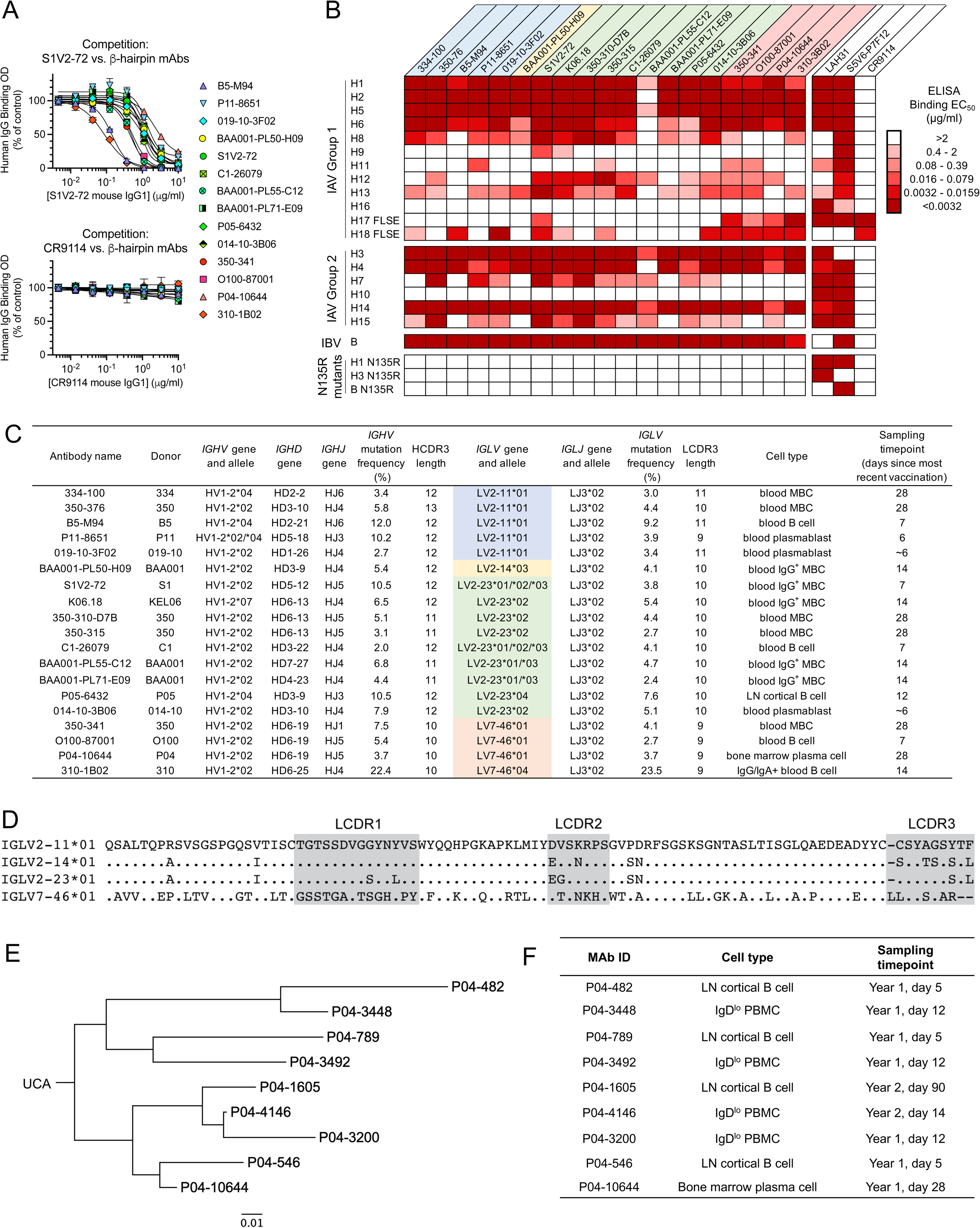
Features of human B-cell receptors that bind the HA2 β-hairpin. A) Results from competitive-binding ELISAs. Binding of putative β-hairpin mAbs (human IgG1) to EHA2 was measured in the presence of serially diluted S1V2-72 mouse IgG1 (top) or CR9114 mouse IgG1 (bottom). Error bars: mean ± S.D. of two replicates. B) A heatmap reporting the binding strength of β-hairpin IgGs for IAV and IBV HAs, measured by ELISA. Each column depicts the results for one recombinant IgG. Column headers are colored according to the *IGLV* gene used by the Ab’s light chain: blue = *IGLV2-11*, yellow = *IGLV2-14*, green = *IGLV2-23*, pink = *IGLV7-46*, white = control IgGs. C) Genetic features and cellular source of each β-hairpin mAb. Abs are colored as in (B). MBC, memory B cell; LN, lymph node. D) Sequence alignment of the polypeptides encoded by the germline human *IGLV* genes commonly used to construct the light chains of β-hairpin Abs. Periods denote unity with the reference sequence, *IGLV2-11*01*; dashes denotes gaps. For simplicity, only one allele is shown per gene. E) Cloanalyst-inferred clonal lineage of Ab P04-10644. F) Cell types and sampling timepoints of the members of the clonal lineage shown in (E), as reported in (*29*).

Each β-hairpin mAb has a HC encoded by *IGHV1-2*, implying that engagement of the β-hairpin epitope requires residues uniquely encoded by this gene (**Fig. 1C**). At least four different LC variable (V) genes permit β-hairpin binding: *IGLV2-11*, *IGLV2-14*, *IGLV2-23*, and *IGLV7-46* **(Fig. 1C)**. The *IGLV2* genes all encode similar amino-acid sequences, from which the sequence of the *IGLV7-46* gene product differs substantially (**Fig. 1D**), raising the possibility of different modes of antigen binding.

B cells that recognize the HA β-hairpin can participate in all the main branches of T cell-dependent humoral responses. The HC and LC V-genes of each β-hairpin mAb have several somatic mutations (**Fig. 1C**), indicating that the corresponding B cells all participated in germinal center reactions. Most of the β-hairpin mAbs were isolated from activated B cells, memory B cells (MBCs), or plasmablasts circulating in peripheral blood, but mAb P05-6432 was from a B cell taken from the cortex of a draining lymph node, and mAb P04-10644 was from a CD138^+^ bone-marrow plasma cell (**Fig. 1C**). Moreover, mAb P04-10644 belonged to a clonal lineage whose nine members were recovered from the draining lymph node, peripheral blood, and bone marrow at various times after seasonal influenza vaccination in two successive years (**Fig. 1E-1F**). Previously reported Ig-Seq serum proteomics analysis showed that Abs matching the B5-M94 or C1-26079 clonotypes contributed 0.7-2.1% of the day 28 serum IgG response specific to H3 A/Singapore/INFIMH-16–0019/2016 HA, as quantified by LC-MS/MS extracted ion chromatogram (XIC) peak area, in two donors who had received the 2018-2019 quadrivalent egg-based (B5-M94) or cell-based (C1-26079) seasonal flu vaccine (*24*). These two clonotypes were also detected in each donor’s serum immediately before vaccination, albeit at lower abundance (*24*), suggesting that β-hairpin clonotypes can be recalled and boosted by seasonal vaccination. We conclude that in many donors, β-hairpin-specific B cells can in principle contribute to influenza immunity as circulating memory B cells and/or as Ig-secreting, bone marrow-resident plasma cells, after undergoing possible affinity maturation in germinal centers.

### Structure of S1V2-72 Fab bound to postfusion HA2

Our previous cryo-EM analysis did not resolve the specific atomic contacts between any β-hairpin mAb and HA2 (*19*). We prepared new cryo-EM maps of S1V2-72 Fab bound to EHA2 from H3 A/Aichi/2/1968 X-31 and B/Malaysia/2506/2004, at respective nominal resolutions of Å and 3.0 Å (**Fig. S2, S3**). Atomic models fit to these maps showed that each Fab molecule interacts with a single protomer of HA2 (**Fig. 2A, Fig. S3G**).

**FIGURE 2.**
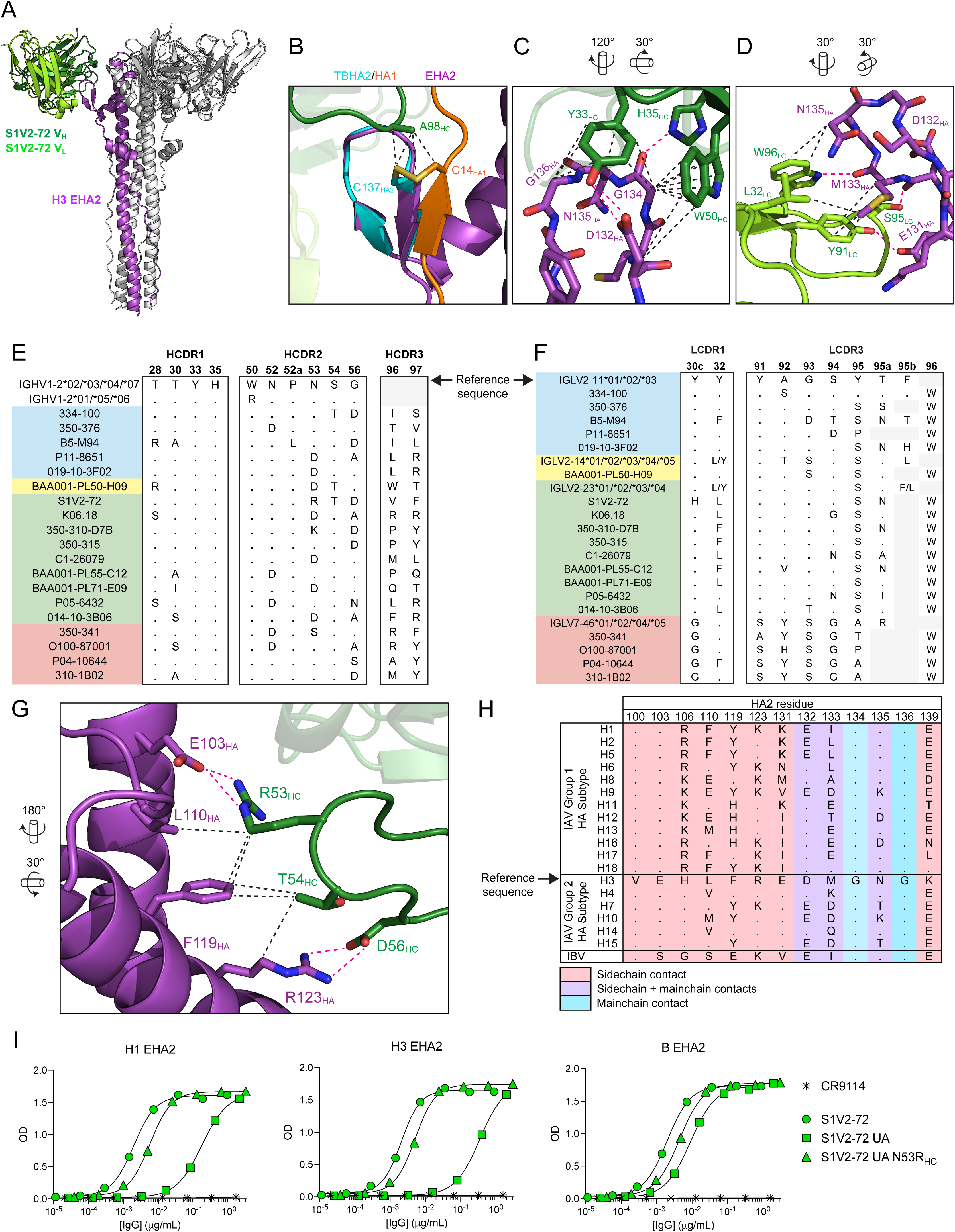
Structure of S1V2-72 Fab bound to postfusion HA2. A) Model of S1V2-72 Fab bound to H3 EHA2. Only one protomer of the Fab:EHA2 complex is colored. Subsequent panels indicate the rotations applied to the view in (A) to achieve the views shown in those panels. B) Structure of the S1V2-72 Fab:EHA2 complex superposed on the crystal structure of H3 TBHA2+HA1 (PDB: 1HTM). Black dashed lines indicate non-polar interactions that might occur in a hypothetical complex of S1V2-72 with bona fide, postfusion HA2+HA1. C) Interactions of S1V2-72 HC with the H3 HA2 β-hairpin. Black dashed lines, nonpolar interactions; magenta dashed lines, polar interactions. D) Interactions of S1V2-72 LC with the H3 HA2 β-hairpin, denoted as in (C). E) Sequence alignment comparing selected residues from β-hairpin Ab HCs and from the polypeptides encoded by germline *IGHV1-2* alleles. F) Sequence alignment comparing selected residues from β-hairpin Ab LCs and from the polypeptides encoded by germline *IGLV* genes. G) Interactions of S1V2-72 HC with the ascending and descending α-helices of H3 EHA2. H) Sequence alignment of the S1V2-72 epitope residues (from the S1V2-72 Fab:H3 EHA2 complex) with the same residues from other IAV and IBV HA subtypes. I) ELISA binding curves for S1V2-72 IgGs. UA, unmutated ancestor. In (C), (D), and (G), parts of the model are hidden for clarity. In (E) and (F), gray spaces denote residues absent from the sequences.

In both structures, the tip of the β-hairpin lies between the tips of the HC complementarity-determining region 1 (HCDR1), HCDR2 and HCDR3 loops; the hairpin also contacts LCDR1 and LCDR3. In authentic HA2, the hairpin has a disulfide link to the N-terminal strand of HA1. Aligning the coordinates for the hairpin plus the disulfide-bonded strand from postfusion TBHA (*31*) with our models showed that the HA1 strand would not necessarily proscribe the Fab:HA2 contact. Instead, the HA1 strand might strengthen Fab binding through van der Waals contacts with the disulfide or other stabilizing contacts with HA1 (**Fig. 2B**). Nevertheless, adding the HA1 strand to the Ab:HA2 complex would require accommodation to avoid steric clashes, by one of at least two potential ways. Either the HA2 hairpin might bend further away from the coiled-coil than in the current structures, so that the adjoining HA1 strand would not clash with side chains projecting from the coiled-coil; or else in the antibody-bound state, the N- and C-terminal regions of the HA1 strand might divert from the antibody, so that only a few residues around C14_HA1_ would make contact with it.

In the structures of S1V2-72 complexed with H3 or B EHA2, several germline-encoded residues from both HC and LC make polar or nonpolar contacts with the conserved HA2 β-hairpin. The side chains of Y33_HC_, H35_HC_, and W50_HC_ contact main-chain atoms in the loop of the HA2 β-hairpin (G134_HA2_, N135_HA2_, G136_HA2_) (**Fig. 2C, Fig S4B**). Y33_HC_ and W50_HC_ also interact with the conserved side-chain of D/E132_HA2_ (**Fig. 2C, Fig S4B**). Similarly, the sidechains of LC germline residues L32_LC_, Y91_LC_, S95_LC_ and W96_LC_ form hydrogen bonds or nonpolar interactions with main-chain atoms and/or conserved side chains of 131_HA2_, 132_HA2_, 133_HA2_, 135_HA2_, and 139_HA2_ (**Fig. 2D, Fig S4C**). The invariance of paratope residues Y33_HC_, H35_HC_, W50_HC_, and W96_LC_ in all β-hairpin mAbs (**Fig. 2E-2F**) suggests that 1) these residues are critical for HA2 binding, and 2) the interactions modeled between these residues and the S1V2-72 epitope apply to complexes of any β-hairpin Ab bound to an EHA2. Likewise, Y91_LC_ is conserved in every LC derived from *IGLV2-11, IGLV2-14,* or *IGLV2-23* (but not those from *IGLV7-46*)(**Fig. 2F**); thus, we surmise that in the corresponding Abs Y91_LC_ makes reproducible contacts crucial for HA2 binding.

In addition to binding the HA2 β-hairpin, the S1V2-72 HC contacts several polar and non-polar residues in the ascending and descending α-helices of HA2 (**Fig 2G, Fig S4D**). Many, but not all, of these contacts involve conserved epitope residues (**Fig 2H, Fig S4E**), which probably explains how S1V2-72 binds so many subtypes of HA. The corresponding paratope residues are also conserved among many β-hairpin Abs, (**Fig. 2E**), suggesting that other β-hairpin Abs may contact the HA2 α-helices similarly.

A few paratope-epitope contacts merit further discussion:

1) The Y91_LC_ hydroxyl group can donate a hydrogen bond either to the carboxylate of E131_HA2_ (**Fig 2D**), which is conserved in Group 2 HA subtypes (**Fig 2H**), or to the carboxylate of E139_HA2_ (**Fig S4C**), which is conserved in IBV and 12/18 IAV subtypes (**Fig 2H**). Only H11, H16, and H17 lack carboxylate sidechains at both 131_HA2_ and 139_HA2_ (**Fig 2H**), which may explain why many β-hairpin Abs do not bind these subtypes (**Fig 1B**).

2) At the tip of the LCDR3 loop, the hydroxyl group of S95_LC_ donates a hydrogen-bond to the carbonyl group of 132_HA2_ (**Fig 2D, Fig S4C**). In 9-residue LCDR3s, the analogous position would be residue 94; in 11-residue LCDR3s, it would be residue 95a. Among all the *IGLV2*-family LCs, 80% (12/15) have Ser/Thr at this position (**Fig 2F**).

3) R53_HC_ makes salt-bridges with E103_HA2_ (conserved in all IAV subtypes)(**Fig 2G-2H)** or with E119_HA2_ in IBV **(Fig S4D**). The alkyl portion of the R53_HC_ side chain also contributes to nonpolar contacts with hydrophobic HA2 residues in both IAV and IBV. The germline residue, N53_HC_, would not make such favorable contacts. Indeed, in ELISAs, an N53R_HC_ substitution increased the avidity of the S1V2-72 unmutated ancestor (UA; i.e., germline-reverted S1V2-72) IgG for H1 and H3 EHA2s by ≥25-fold (**Fig. 2I**). No other β-hairpin Ab has R53_HC_, but 350-310-D7B has K53_HC_ (**Fig 2E**), which may recapitulate some of the favorable contacts of R53_HC_. The ARMADILLO model (*32*) of Ab somatic mutation predicts that the N53R_HC_ substitution is rare: in a pool of *IGHV1-2*-derived HCs with an average V_H_ mutation frequency of 10%, only ∼1.4% of the sequences would be expected to have R53_HC_, while ∼5.5% would be expected to have K53_HC_. Thus, the N53R_HC_ substitution appears to be an improbable-but-significant step in the affinity maturation of S1V2-72. Nevertheless, the N53R_HC_ mutation is not the only pathway for achieving tight binding to H1, H3, and B HAs, as the other β-hairpin Abs demonstrate.

In addition to binding postfusion HA2, S1V2-72 can bind solubilized, full-length HA ectodomain in its prefusion conformation, albeit with lower avidity (*19*). It is unclear whether recognition of prefusion or postfusion HA is what allows passively transferred S1V2-72 IgG2c to mediate protection from severe influenza disease in lethally challenged mice (*19*). Alignment of the HA2 β-hairpin from the S1V2-72:H3 EHA2 complex with that from a crystal structure of prefusion HA0 shows that several residues included in the postfusion epitope would be incorrectly positioned for S1V2-72 binding to prefusion HA (**Fig S4F**). Moreover, partial occlusion of the β-hairpin in the prefusion HA homotrimer would cause a severe steric clash with the S1V2-72 HC (**Fig S4G**). Prefusion HA may explore conformations other than the canonical, steady-state form observed in x-ray crystallography and cryo-EM studies; we and others have proposed that these alternate conformations transiently expose otherwise occluded epitopes (*18, 33*). Such molecular “breathing” may allow β-hairpin Abs to recognize prefusion HA on the surface of flu-infected cells. Experiments to distinguish these possibilities are underway.

### Structure of 350-341 Fab bound to H3 EHA2

The lack of conservation between the LCDR3s of *IGLV2* and *IGLV7-46* LCs – particularly the lack of Y91_LC_ in the latter (**Fig. 2F**) – implies that these V-genes offer different solutions for binding the β-hairpin. We prepared a cryo-EM map of a representative *IGLV7-46*-derived β-hairpin mAb, 350-341, bound to EHA2 from H3 A/Aichi/2/1968 X-31 (**Fig. S5**). The atomic model fit to this 3.5Å-resolution map showed that 350-341 binds HA2 in a manner grossly similar to S1V2-72 (**Fig. 3A**), but the degree to which the HA2 β-hairpin bends away from the coiled-coil varies between the two structures (**Fig. 3B**). In the 350-341:HA2 complex, the side chains of germline-encoded residues Y34_LC_, D50_LC_, and S93_LC_, and the Cα of G94_LC_ make important contacts with the β-hairpin (**Fig. 3C**); these contacts are not recapitulated in the S1V2-72:EHA2 complex. The paratope residues of the 350-341 LC are generally conserved in the other *IGLV7-46* Abs (**Fig. 3D**), suggesting that their LCs bind HA2 in the same fashion. We conclude that β-hairpin Abs comprise two public sub-classes distinguished by their LCs. The first sub-class (“type 1”), exemplified by S1V2-72, uses *IGLV2*-family genes to construct its LCs. The second sub-class (“type 2”), typified by 350-341, uses *IGLV7-46*. Although the *IGLV7-46* and *IGLV2* LCs are structurally distinct, they all use *IGLJ3*02-*encoded W96_LC_ to bind the backbone carbonyl of 133_HA2_ (**Fig. 2D, 2F; Fig. 3C, 3D**).

**FIGURE 3.**
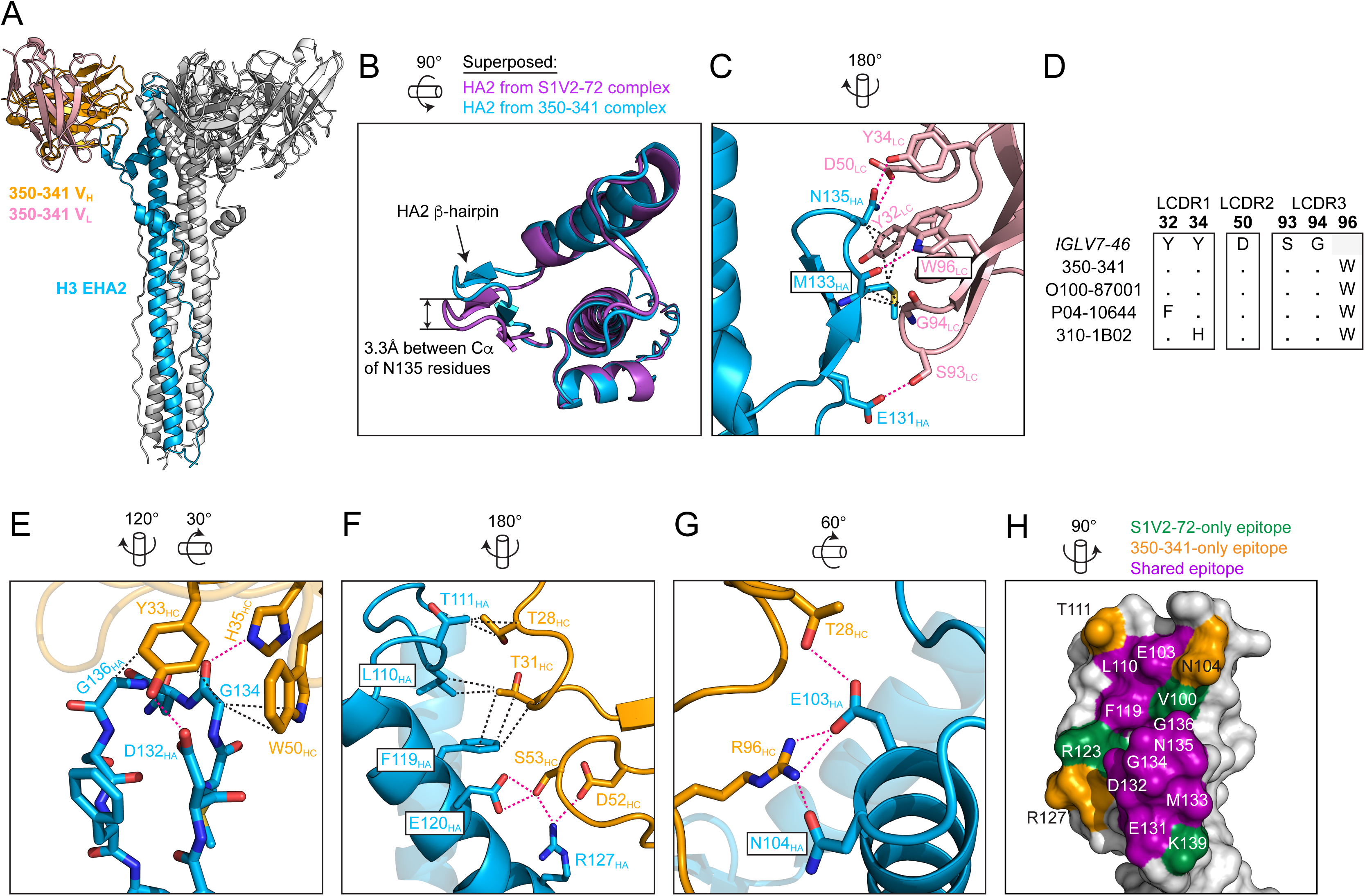
Structure of the 350-341 Fab bound to postfusion HA2. A) Model of 350-341 Fab bound to H3 EHA2. Only one protomer of the Fab:EHA2 complex is colored. Subsequent panels indicate the rotations applied to the view in (A) to achieve the views shown in those panels. B) Top view of the superposed structures of the H3 EHA2s from the S1V2-72:EHA2 complex and 350-341:EHA2 complex. C) Interactions of 350-341 LC with the H3 HA2 β-hairpin. Black dashed lines indicate nonpolar interactions; magenta dashed lines indicate polar interactions. D) Sequence alignment comparing selected residues from the polypeptide product of *IGLV7-46* and the same residues from HA2 β-hairpin Abs that use IGLV7-46 LCs. Dots denote unity with the germline sequence. E) Interactions of 350-341 HC with the H3 HA2 β-hairpin, denoted as in (C). F) Interactions of 350-341 HC with the ascending and descending α-helices of H3 EHA2. G) Additional interactions of 350-341 HC with α-helices of H3 EHA2. H) Surface rendering of H3 EHA2 showing the overlap of the S1V2-72 and 350-341 epitopes. In (B), (C), and (E-H), parts of the model are hidden for clarity.

Like S1V2-72, the 350-341 HC contacts the HA2 β-hairpin through *IGHV1-2*-encoded Y33_HC_, H35_HC_, and W50_HC_ residues (**Fig. 3E**). The 350-341 HC also makes polar and nonpolar interactions with conserved HA2 residues outside the β-hairpin (**Fig. 3F-3G**). Some of the epitope contacts – including the sidechains of E103_HA2_, L110_HA2_, and F119_HA2_ – are shared by 350-341 and S1V2-72 (**Fig. 3H**). The corresponding paratope residues are not conserved between the two Abs, however. For example, the guanidinium group of R96_HC_ in 350-341 binds the carboxylate of E103_HA2_ (**Fig. 3G**), while R53_HC_ in S1V2-72 makes this contact instead (**Fig. 2G**). Other epitope residues are bound uniquely by 350-341 or S1V2-72 (**Fig 3H**). Thus, although 350-341 and S1V2-72 belong to the same class of *IGHV1-2*-derived β-hairpin Abs, their HCs arrive at somewhat different solutions for binding the region around the β-hairpin.

### Genetic restrictions for germline HCs of β-hairpin Abs

Known β-hairpin Abs all derive from *IGHV1-2* (**Fig. 1C**), raising the question of whether any other germline *IGHV* gene encodes the residues necessary to recognize the β-hairpin. We determined the germline residues critical for β-hairpin binding by generating a series of recombinant IgGs in which individual paratope residues from the S1V2-72 UA (i.e., the germline antigen receptor that would be expressed by a naïve B cell) were converted to Ala. ELISA analysis of the mutant IgGs showed that *IGHV*-encoded Y33_HC_, H35_HC_, and W50_HC_ are independently crucial for the S1V2-72 UA to bind H1, H3, and B EHA2 (**Fig. 4A**). These residues are critical probably because their sidechains make several contacts with main-chain atoms and conserved sidechains in the β-hairpin. Only two *IGHV* genes encode the complete triad of Y33_HC_, H35_HC_, and W50_HC_: *IGHV1-2* (alleles *\*02, *03, *04*, and *\*07*) and *IGHV1-45* (*\*01, *02, *03*, i.e., all alleles) (**Fig. 4B**). We have not found any β-hairpin Ab constructed from *IGHV1-2*01/*05/*06*, which encode Y33_HC_ and H35_HC_, but use R50_HC_ instead of W50_HC_; thus, we tentatively surmise that R50_HC_ is incompatible with β-hairpin binding. We cannot exclude the possibility that other amino acids could functionally substitute for Y33_HC_, H35_HC_, or W50_HC_, which might enlarge the pool of *IGHV* genes capable of encoding a permissive HC. Nevertheless, the absolute conservation of these three residues in all known β-hairpin Abs (**Fig. 2E**) implies that Y33_HC_, H35_HC_, and W50_HC_ are the favored – if not only – amino acids for binding the HA2 β-hairpin.

**FIGURE 4.**
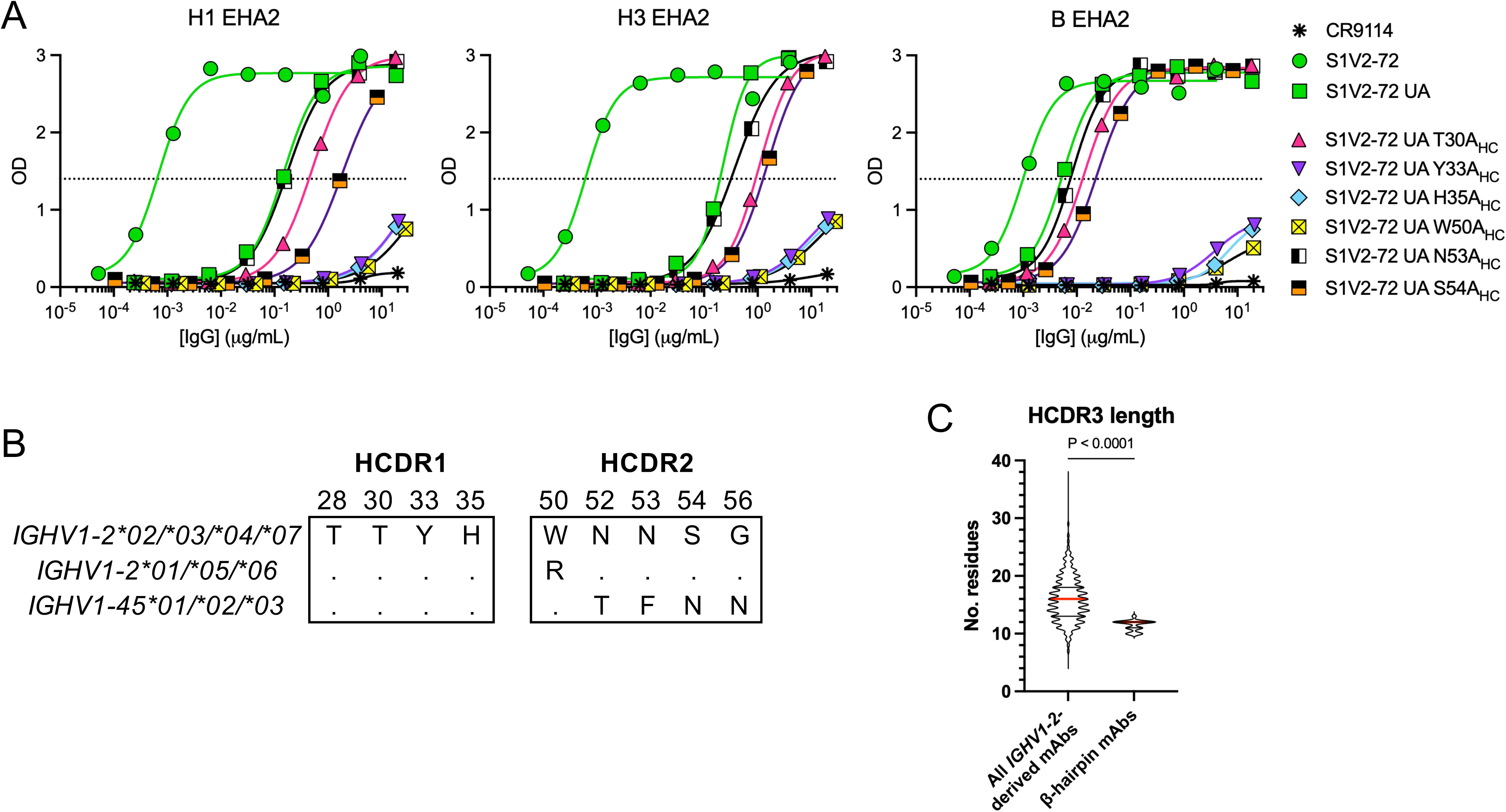
Genetic restrictions for germline forms of β-hairpin Ab HCs. A) Binding of S1V2-72 unmutated ancestor (UA) IgGs to EHA2s, as measured by ELISA. B) Sequence alignment showing conservation of key paratope residues in *IGHV1-2*\**02/*03/*04/*07* and *IGHV1-45*. C) Violin plot depicting the distribution of HCDR3 lengths for all *IGHV1-2*-derived HCs (n=12,073) from donors P04+P05+P11 or for the *IGHV1-2*-derived HCs from β-hairpin Abs (n=19). The means of the two groups were compared with Welch’s t-test for unequal variance (two-tailed). Thick, red, horizontal lines denote the medians; thin, black, horizontal lines denote the upper and lower quartiles.

Another apparent constraint for β-hairpin Abs is their HCDR3 length. The mean HCDR3 length for all 12,073 *IGHV1-2*-derived HCs recovered from donors P04+P05+P11 is 16.0 ± 3.8 residues (mean ± S.D.)(**Fig. 4C**). In contrast, β-hairpin Abs have shorter HCDR3s (11.4 ± 0.9 residues)(**Fig. 4C**), probably to avoid clashes with the HA1 strand that would be tethered to the HA2 β-hairpin in forms of HA present during viral infection or seasonal influenza vaccination (**Fig. 2B**).

### Genetic constraints for germline LCs of type 1 β-hairpin Abs

The LCs of type 1 β-hairpin Abs use *IGLV2-11, IGLV2-14,* or *IGLV2-2*3 joined to *IGLJ3*02* (**Fig. 1C**). To understand why these genes are favored for LC construction, we defined the critical determinants of HA2 binding, using the S1V2-72 UA LC as a model. ELISA analysis of IgGs in which the paratope residues of the germline LC were individually mutated to Ala showed that V-gene-encoded Y91_LC_ and J-gene-encoded W96_LC_ are critical for recognition of H1, H3, and B EHA2 (**Fig. 5A**). Twenty-four functional *IGKV* and *IGLV* genes encode the critical Y91 (**Fig. 5B**), while *IGKJ1*01* and *IGLJ3*02* encode the necessary W96.

**FIGURE 5.**
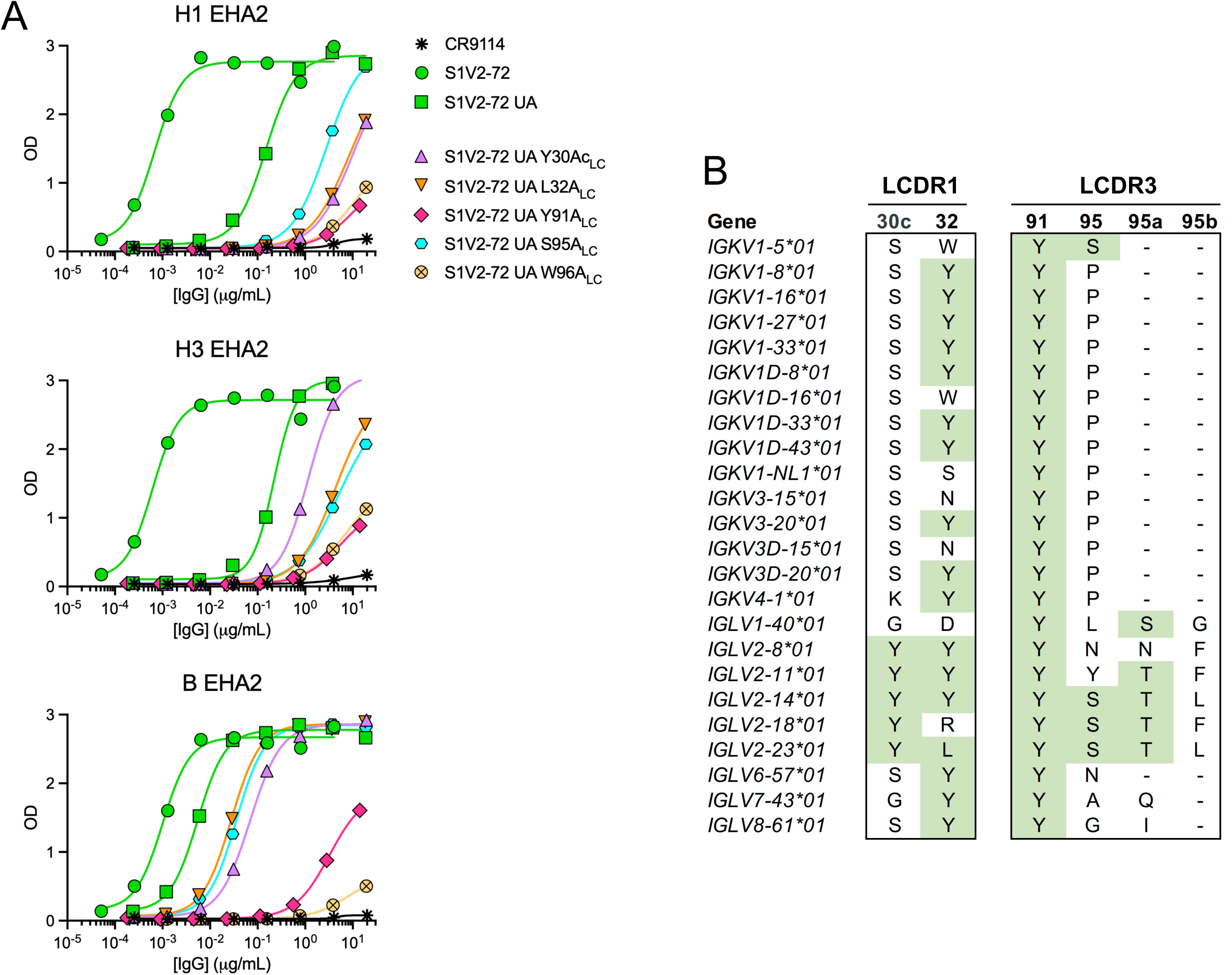
Genetic constraints for germline LCs of type 1 β-hairpin Abs. A) Binding of S1V2-72 unmutated ancestor (UA) IgGs to EHA2s, as measured by ELISA. B) Sequence alignment comparing the residues encoded by selected LC V-genes at positions that contribute to the type I LC paratope.

Germline residues L32_LC_ and S95_LC_ also contribute appreciably to S1V2-72’s affinity for EHA2 (**Fig. 5A**). S95_LC_ immediately precedes the V-J junction and therefore does not have to be germline-encoded, since its codon could be supplied by exonuclease activity plus palindromic (P-) and nontemplated (N-) nucleotide addition during V_L_-J_L_ recombination. However, because nucleotide trimming and nontemplated addition are generally minimal in human V_L_-J_L_ rearrangement (*34–37*), construction of type 1 LCs likely favors V-genes that already encode S95(a) or the functionally similar T95(a). This prediction is supported by the exclusive use of *IGLJ3*02* to encode W96_LC_ on the C-terminal flank of the V-J junction (**Fig. 1C**). Only six of the 24 V-genes that encode Y91 also have S/T95(a): *IGKV1-5*, *IGLV1-40*, *IGLV2-11, IGLV2-14, IGLV2-18*, and *IGLV2-2*3. Of these six genes, only *IGLV2-11, IGLV2-14,* and *IGLV2-23* encode L32_LC_ or the Y32_LC_ substitution present in many type I LCs (**Fig. 5B**).

### Limitations to the binding breadth of β-hairpin mAbs

β-hairpin mAbs bind weakly – if at all – to H9 and H10 (**Fig. 1A**), the only subtypes to have K135_HA2_ (**Fig. 2H**). Mutation of K135_HA2_ to the consensus residue, Asn (K135N_HA2_), substantially tightened binding of several β-hairpin mAbs to H9 and H10 (**Fig. 6A, Fig. S6A**-**S6B**). The inability of β-hairpin Abs to recognize HA2s with either K135_HA2_ or R135_HA2_ (**Fig 1B**) suggests that bulky and/or positively charged sidechains at position 135_HA2_ generally proscribe binding. Some IgGs were unable to bind even the K135N_HA2_ versions of H9 and H10 (**Fig. 6A**, **Fig S6A-S6B**), implicating secondary impediments to binding HAs of these subtypes.

**FIGURE 6.**
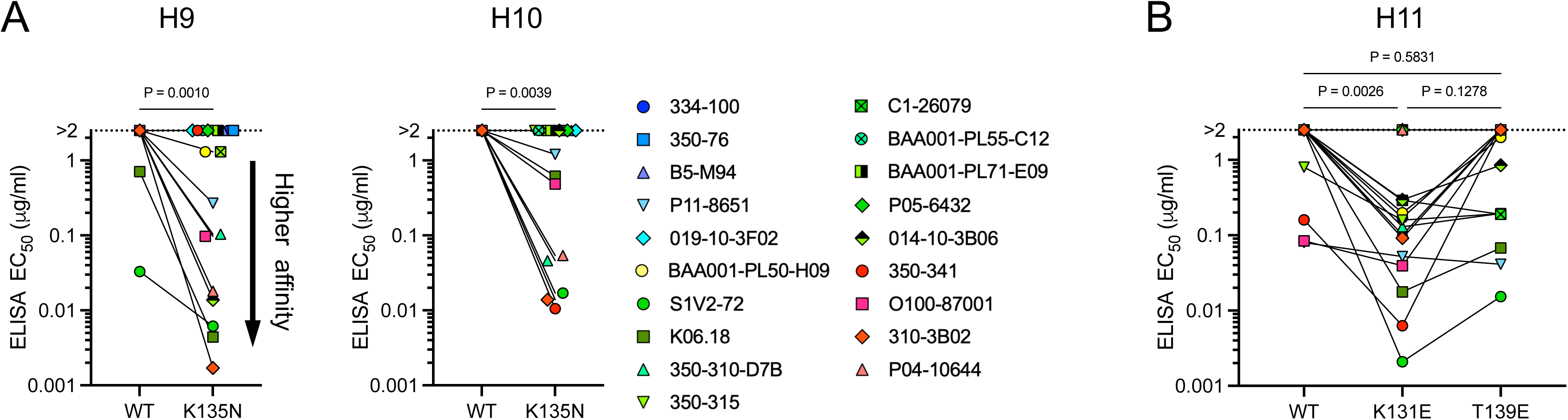
HA2-intrinsic impediments to recognition of H9, H10, and H11 by β-hairpin Abs. A) ELISA EC_50_ values for IgG binding to wildtype and K135N_HA2_ EHA2s. B) ELISA EC_50_ values for IgG binding to wildtype, K131E_HA2_, or T139E_HA2_ H11 EHA2. The statistical analyses used were a one-tailed, Wilcoxon matched-pairs signed rank test (A) or the Friedman test (P=0.0002) with Dunn’s post test, corrected for multiple comparisons (B). The P-values from Dunn’s post test are reported (B).

Most type 1 β-hairpin mAbs also fail to bind H11, H16, and H17 (**Fig. 1A**), probably due at least in part to the lack of favorable interaction with the side chains of 131_HA2_ or 139_HA2_. Our cryo-EM structures show that S1V2-72 and probably all type 1 β-hairpin Abs donate a hydrogen bond from the hydroxyl group of Y91_LC_ to the carboxylate moiety of either 131_HA2_ or 139_HA2_, found in most HA subtypes (**Fig. 2D, 2H, S4C**). Only H11, H16, and H17 lack carboxylates at both positions (**Fig 2H**). Conversion of either K131_HA2_ or T139_HA2_ to Glu allowed S1V2-72 and several other type 1 β-hairpin Abs to bind H11, with E131_HA2_ generally conferring tighter binding than E139_HA2_ (**Fig. 6B, Fig S6C**). However, neither I131E_HA2_ nor N139E_HA2_ mutations permitted binding to H16 (data not shown). We did not test whether analogous mutations rescue binding to H17.

Half of the type 2 β-hairpin mAbs also fail to bind H11, and none bind H16 (**Fig 1A**). The structure of 350-341 bound to H3 EHA2 shows that type 2 LCs can donate a hydrogen bond from the hydroxyl group of S93_LC_ to the carboxylate of E131_HA2_, which is conserved in all IAV group 2 subtypes. It is plausible that S93_LC_ might accept a hydrogen bond from the ε-amino group of K131_HA2_ in H1, H2, H5, and H11, but E131_HA2_ supplies a more favorable interaction, as shown by the tighter binding of 75% of type 2 Abs to H11 with an K131E_HA2_ mutation (**Fig. 6B**). Conversion of the nearby T139_HA2_ to Glu eliminates binding of type 2 Abs to H11 (**Fig 6B**); we speculate that E139_HA2_ sequesters the charged sidechain of K131_HA2_ away from S93_LC_. As with type 1 Abs, neither I131E_HA2_ nor N139E_HA2_ mutations permitted type 2 Abs to bind H16 (data not shown).

## DISCUSSION

Antigenic drift in response to population-level immune pressure is a persistent issue for human immunity to influenza viruses, necessitating yearly updates to the seasonal flu vaccine as the viruses accrue escape mutations. A vaccine that elicits protective Abs to broadly conserved flu epitopes would not require annual reformulation and would protect against pre-pandemic strains in the event of spillover into humans. Here, we analyze a public class of protective human antibodies (*19*) that recognize an epitope encompassing a β-hairpin and adjacent residues exposed on the surface of postfusion HA2. This β-hairpin epitope is broadly conserved among seasonal and pre-pandemic flu viruses, including in highly pathogenic H5 and H7 strains that have caused severe disease in humans (*6, 19*).

The β-hairpin epitope may owe its conservation to functional constraints or to a lack of population-level immune pressure. Excluding the former possibility is non-trivial because there is no established, *in vitro* method for selecting escape mutations from non-neutralizing Abs. As for immune pressure, we previously estimated from two donors that ≤0.1% of circulating, HA-binding memory B cells target the β-hairpin (*19*), and, as we report here, clonotypes matching β-hairpin Abs B5-M94 or C1-26079 constituted ≤2.1% of the anti-H3 serum IgG response in two additional donors. This lack of immune pressure may extend to the wider population. If so, then raising the salience of the HA2 β-hairpin by epitope-targeted vaccination could trigger a cyclical “arms race” between mutating virus and mutating B cells. Our findings here might forecast potential opening moves of such a competition: viral acquisition of K/R135_HA2_ is a possible escape route from many β-hairpin Abs, as is simultaneous elimination of carboxylate side chains at both 131_HA2_ and 139_HA2_. Potential counter-moves are foreshadowed by S1V2-72, 350-341, and other β-hairpin Abs that still bind HAs with these escape mutations. Regardless of any functional constraints on the epitope or capacity of humoral immunity to counter viral mutations, the strongest vaccine design may be one that targets a combination of spatially separated, conserved epitopes, e.g., the β-hairpin plus the receptor binding site (*16, 17*), head interface (*18, 38*), central stem (*15, 39*), etc. Such an approach would raise the barrier to viral escape by necessitating improbable, simultaneous mutations at several epitopes.

It will be important to resolve whether β-hairpin Abs mediate protection from severe disease by recognizing the prefusion or postfusion conformation of HA. Although the β-hairpin is conformationally conserved between pre- and postfusion states, the surrounding residues that contribute to the S1V2-72 epitope are quite different in pre- and postfusion HA. Presumably, the ideal immunogen for eliciting potently protective β-hairpin Abs is whichever HA conformation the Abs bind on the surface of infected cells. If the postfusion conformation is the relevant in vivo state, it will be necessary for the postfusion HA2 immunogen to include at least a fragment of the HA1 strand that in native HA1+HA2 remains tethered by disulfide linkage to C137_HA2_. Without this linked HA1 strand in place to enforce selection for Abs with HCDR3s ≤13 residues, many of the Abs elicited by vaccination with postfusion HA2 will probably have longer HCDR3s – as is typical for human Ab HCs – and would therefore be unable to bind HA1+HA2 on flu-infected cells.

Accumulating evidence for the protective role of non-neutralizing Abs during influenza infections indicates that β-hairpin Abs could confer important protection from severe influenza disease. Levels of serum Abs to the HA stem are a correlate of protection from disease progression after natural infection with H1N1 influenza (*40–42*). Lifelong protection against severe disease and death after infection by highly pathogenic H5N1 or H7N9 virus has also been attributed to cross-reactive stem Abs elicited by childhood infection with seasonal flu strains (*43*). Several studies have shown that these HA stem Abs confer protection through Fc receptor-dependent mechanisms, unless the Abs are artificially infused to serum and mucosal concentrations higher than those likely to be elicited by vaccination or infection, at which point the Abs also may have neutralizing activity (*20–22, 42, 44, 45*). Therefore, β-hairpin Abs, which protect mice from lethal IAV and IBV challenge via Fc effector functions (*19*), may also help protect humans from severe influenza.

Isolation of HA2 β-hairpin mAbs from 14 genetically unrelated donors suggests that HA exposure routinely raises these humoral responses in humans. But what fraction of the human population has the immunoglobulin genes needed to make β-hairpin Abs? Probably all humans can assemble an appropriate LC, because there are several *IGLV* genes that permit binding to the β-hairpin, and the critical W96_LC_ residue can be encoded either by *IGLJ3*02* or by P- or N-nucleotide addition during V-J recombination. As for the HC, both *IGHV1-2*02/*03/*04/*07* and all alleles of *IGHV1-45* encode a triad of paratope residues (Y33_HC_, H35_HC_, W50_HC_) favorable for binding the β-hairpin. We have not yet isolated any *IGHV1-45*-derived β-hairpin Abs from humans, however. If, for whatever reason, *IGHV1-45* cannot contribute to a β-hairpin Ab HC, then >75% of people with ethnic origins in South Asia or East Asia have at least one *IGHV1-2* allele permissive for β-hairpin binding, while >85% of individuals with ethnic origins in Africa or the Americas and >90% of those with European ancestry have at least one permissive *IGHV1-2* allele (*46*). We cannot exclude that other *IGHV* genes may be permissive for β-hairpin binding without using Y33_HC_+H35_HC_+W50_HC_, especially if the resultant HCs bind the β-hairpin with a different footprint than the examples we have described here.

Some broadly protective Abs to influenza and HIV-1 must follow tortuous, specific paths of affinity maturation to evade immune tolerance controls or to acquire binding breadth and potency (*47, 48*). Indeed, the germline ancestors of certain broadly protective HIV-1 Ab lineages bind the desired antigen so weakly that regimens of specially designed, germline-targeting immunogens are needed to coax the B cells toward their ultimate, broadly protective destiny (*49*). Such measures are probably unnecessary for eliciting HA2 β-hairpin Abs. Breadth of binding to H1+H3+B EHA2s appears to be an inherent, germline property of some, if not many, HA2 β-hairpin Abs, including S1V2-72. Substantial avidity for HA2 is also a common germline property of these Abs. Further acquisition of breadth and potency does not necessarily require extensive somatic mutation, for a single mutation (N53R_HC_) to germline S1V2-72 suffices to increase its divalent avidity for H1 and H3 by >25-fold. Nor must β-hairpin Abs follow a single path to attain potency, because most β-hairpin Abs achieve high avidity for H1, H3, B and other HA subtypes without the N53R_HC_ mutation. Finally, that ≥14 donors produce anti-β-hairpin humoral responses suggests that tolerance controls do not routinely proscribe these responses. Therefore, elicitation of potent, broadly protective β-hairpin humoral responses may be a relatively straightforward task for vaccinologists.

### Limitations of the study

We identified several of the β-hairpin mAbs by searching published datasets for Abs that matched a specific genetic signature (i.e., a HC encoded by *IGHV1-2* plus a LC encoded by *IGLV2* genes or *IGLV7-46*). Our search method ignored any potential β-hairpin Abs that did not match this genetic signature. Other types of β-hairpin Abs, should they exist, could be identified by sorting EHA2-binding B cells directly from donor samples, or by other methods. In native HA1+HA2, a strand of HA1 forms a disulfide linkage to the HA2 β-hairpin, but the Fab:HA2 complexes we studied here lack the HA1 subunit. It will be important to resolve the extent to which the HA1 subunit changes the footprint of β-hairpin Abs and impacts the stability of the Fab:HA complex.

## ACKNOWLEDGMENT

We thank the Harrison laboratory and the many members of the Program Project consortium for structure-function analysis of influenza virus infection- and vaccine-induced B-cell repertoires for advice and discussion. We are grateful for the excellent technical support of R. Abbott, K. Boradia, L. Zhu, and J. Li. This research was conducted in part using equipment and services provided by the Harvard Cryo-EM Center for Structural Biology. Some molecular graphics were rendered with UCSF ChimeraX, developed by the Resource for Biocomputing, Visualization, and Informatics at the University of California, San Francisco, with support from National Institutes of Health R01-GM129325 and the Office of Cyber Infrastructure and Computational Biology, National Institute of Allergy and Infectious Diseases. We acknowledge support of grants and contracts from the National Institute Of Allergy And Infectious Diseases of the National Institutes of Health: K99-AI193241 (to JF), P01-AI089618 (to SCH, GK, and MAM), contract 75N93019C00052 (to MAM, sub-contract to GG), and 75N93024C00074 (to GK and MK). The content is solely the responsibility of the authors and does not necessarily represent the official views of the National Institutes of Health. JF is supported by a Career Development Fellowship Award from the Basic & Clinical Translational Research Executive Committees of the Office of Faculty Development of Boston Children’s Hospital. SCH is supported by the Howard Hughes Medical Institute.

## ACCESSION

The nucleotide sequences of the variable domains of the mAbs used in this study have been deposited in GenBank (accession numbers PX910701-PX910744). Cryo-EM maps were deposited at the Electron Microscopy Data Bank (accession numbers EMD-76704, EMD-76710, EMD-76712). Coordinates for the atomic models were deposited at the Protein Data Bank (accession numbers 12RB, 12RK, 12RO).

## MATERIALS AND METHODS

### Study approvals and volunteers

The study procedures, informed consent, and data collection documents were reviewed and approved by the Duke Health Institutional Review Board (Pro00020561, initial approval 2010). Written informed consent was obtained from donor BAA001. Donor BAA001 (born 1994) was administered the 2019-2020 quadrivalent inactivated influenza vaccine, and blood was collected 14 d post-vaccination. Blood samples were processed into PBMCs and plasma and then aliquoted and stored in liquid nitrogen vapor phase or at −80°C, respectively, for future analysis.

### Identification of potential β-hairpin mAbs

We previously reported that HA2 β-hairpin Abs use HCs that are encoded by *IGHV1-2*, and LCs that are encoded by *IGLV2-11* or *IGLV2-23* (*19*). After determining that mAb O100-87001 (Towers *et al.*, in preparation) also binds the HA2 β-hairpin, we expanded the search criteria for potential β-hairpin Abs to include those with 1) a HC encoded by *IGHV1-2*, and 2) a LC encoded by an *IGLV2*-family gene or *IGLV7-46*. We used these criteria to search published paired-chain V(D)J sequence datasets (*25–29*) and/or Ig-Seq serum proteomics datasets (*24*) from donors recently immunized with influenza vaccines. We expressed a subset of the Abs that matched the search criteria as recombinant IgGs for further analysis.

Individual memory B cells of donor BAA001 were sorted into Nojima cultures, and the resultant clonal IgGs were screened for HA-binding essentially as described (*50, 51*). Clonal IgGs that bound H1+H3+B EHA2 and used HCs encoded by *IGHV1-2* were selected for further characterization as recombinant IgGs. A complete account of the experimental methods and results from donor BAA001 is forthcoming (Kelsoe and colleagues, manuscript in preparation).

### V(D)J sequence analysis and clonal lineage inference

Rearranged V(D)J sequences were annotated with IMGT/V-QUEST (*52*). The clonogram for the P04-10644 clonal lineage was inferred with Cloanalyst (*53*) and plotted with FigTree v1.4.5.

### Cell line culture

Expi293F cells (*Homo sapiens*; Thermo Fisher Scientific) were cultured at 37°C, 8% CO_2_, with shaking in a humidified incubator. Expi293F cells were grown in Expi293 Expression Medium plus penicillin (100 units/ml) and streptomycin (100 μg/ml)(all from Gibco). Expi293F cells were not subjected to authentication or mycoplasma testing.

### Recombinant HA expression and purification

FLSEs representing H17 A/little yellow-shouldered bat/Guatemala/164/2009 and H18 A/flat-faced bat/Peru/033/2010 were a gift from the laboratory of Aaron Schmidt.

Except where explicitly noted otherwise in the figures or text, all recombinant HAs were post-fusion HA2 trimers (EHA2s) (*30*). Although “EHA2” originally referred to truncated, *<u>E</u>scherichia coli*-produced <u>HA2</u> lacking the fusion peptide and transmembrane domain (*30*), we continue to use “EHA2” to refer to otherwise identical protein produced in <u>E</u>xpi293F cells.

Synthetic DNA encoding a His_6_-tag, HRV3C protease cleavage site, and EHA2 was cloned into a modified pVRC8400 plasmid (Addgene #63160) between the NheI and NotI sites. Expression plasmid was transiently transfected into Expi293F cells using either the ExpiFectamine 293 transfection kit (Gibco) or polyethylenimine (MilliporeSigma). Supernatant from transfected cells was harvested 5 days post-transfection and clarified by centrifugation. EHA2 was purified by adsorption to Co^2+^–nitrilotriacetic acid (Co-NTA) metal affinity resin (Takara), washed extensively with buffer A (10 mM HEPES, 150 mM sodium chloride, pH 7.5), and eluted with buffer A plus 400 mM imidazole. EHA2 used for cryo-EM was liberated of the His_6_-tag by incubation with His_6_-tagged HRV3C protease (Pierce), according to the manufacturer’s protocol, then His_6_-tagged peptides were removed by another round of adsorption to Co^2+^-NTA resin. EHA2 was exchanged into imidazole-free buffer (typically PBS), concentrated, and stored short-term at 4°C, or flash-frozen and stored long-term at -80°C.

The following EHA2s were thus produced: H1 A/USSR/90/1977, H2 A/Japan/305/1957, H3 A/Aichi/2/1968 X31, H4 A/American black duck/New Brunswick/00464/2010, H5 A/Vietnam/1203/2004, H6 A/chicken/Taiwan/2759/2012 , H7 A/chicken/Guangdong/J1/2017, H8 A/northern shoveler/California/HKWF1204/2007, H9 A/duck/Beijing/MG0617/2005, H10 A/chicken/Jiangxi/18933/2013, H11 A/common teal/Netherlands/1/2015, H12 A/Northern Shoveler/Nevada/D1516557/2015, H13 A/Armenian gull/Republic of Georgia/2/2012, H14 A/blue-winged teal/TX/AI13-1028/2013, H15 A/shearwater/Australia/2576/1979, H16 A/shorebird/Delaware/172/2006, B/Malaysia/2506/2004.

### Recombinant IgG expression and purification

AbVec2.0-IGHG1, AbVec1.1-IGKC, and AbVec2.1-IGLC2-MscI plasmids, which harbor the constant regions of human IgG1, Igκ, or Igλ, and AbVec2.0-mIghg1 and AbVec2.1-mIglc2, which harbor the constant regions of mouse IgG1 or Igλ, were gifts from Hedda Wardemann (RRID:Addgene_80795; RRID:Addgene_80796; RRID:Addgene_80797; RRID:Addgene_127156; RRID:Addgene_127158) (*54*). Synthetic DNAs encoding Ab HC- or LC-variable domains were cloned into these expression vectors. The vectors were transiently transfected into Expi293F cells with the Expifectamine 293 transfection kit (Thermo Fisher), according to the manufacturer’s instructions. Alternatively, polyethylenimine (MilliporeSigma) was used to deliver the plasmids to the cells. Five days post-transfection, supernatants were harvested, clarified by low-speed centrifugation, mixed 1:1 with Protein A binding buffer (human IgGs) or Protein G binding buffer (mouse IgG1), and incubated with Pierce Protein A agarose resin (human IgGs) or Pierce Protein G agarose resin (mouse IgG1) for ≥1 h at 4°C with agitation. The IgG-adsorbed resin was collected in a chromatography column, washed extensively with binding buffer, eluted in Pierce IgG Elution Buffer (Thermo Fisher), neutralized by one-tenth volume of 1M Tris (pH 9), and dialyzed into PBS plus 0.1% sodium azide. IgG concentrations were determined with a NanoDrop spectrophotometer (Thermo Fisher). Purified, concentrated IgGs (≥1 mg/ml) were stored at 4°C until use.

### Recombinant Fab purification

Fab was expressed in Expi293 cells as described for IgGs, except that the AbVec2.0-IGHG1 plasmid was modified to replace the DNA encoding full-length IgG1 constant region with DNA encoding the first constant domain (CH1) of human IgG1 plus a C-terminal HRV3C protease cleavage site and His_6_-tag. His_6_-tagged-Fab was isolated from clarified cell culture supernatant by adsorption to Co-NTA resin, and liberated of the His_6_-tagged-Fab by incubation with HRV3C protease, as described for purification of EHA2s. Purified Fabs were stored at 4°C in PBS plus 0.1% sodium azide.

### ELISA

High-binding 96- or 384-well microplates (Corning, Greiner Bio-One, or Thermo Fisher) were coated with ≥2 μg/ml protein in 0.1 M sodium carbonate buffer (pH 9) overnight at 4°C or for 2 h at room temperature. Coating proteins included goat anti-human Igκ (Southern Biotech), goat anti-human Igλ (Southern Biotech), or rHAs. After washing with PBS plus 0.1% Tween20, the plates typically were blocked with PBS plus 0.5% BSA at RT for ≥30 min or overnight at 4°C. Next, rIgGs diluted in PBS plus 0.5% BSA and 0.1% Tween20 (ELISA dilution buffer) were added to the plates and incubated for 2 h at RT, or overnight at 4°C. After extensive washing, bound Abs were detected with goat anti-human IgG-HRP (Southern Biotech) diluted in ELISA dilution buffer. After washing, HRP activity was detected with the TMB substrate kit (BioLegend) and a Spark plate reader (Tecan). Background signal at 650 nm was subtracted from the signal at 450 nm to calculate the net optical density (OD).

### Competitive inhibition assay

Ab epitope mapping was performed by competitive binding inhibition in ELISA (*19*). ELISA plates were coated with B EHA2, washed, and blocked as described above. Recombinant CR9114 or S1V2-72 mouse IgG1 was serially diluted in ELISA dilution buffer and then incubated with HA-coated plates for 2 h at RT or overnight at 4°C. Recombinant human IgG1s were then added at a fixed concentration to each well, and incubated with the competitor Abs and HA substrate for 2 h at RT. After washing, bound human IgG was detected by mouse anti-human IgG Fc-HRP (JDC-10, SouthernBiotech).

### Cryo-EM sample preparation

For the S1V2-72:H3 complex, H3 EHA2 was mixed with 1.5-fold molar excess (per EHA2 polypeptide chain) of S1V2-72 Fab and diluted to 2.4 mg/ml protein (final) in PBS plus 0.25% (w/v) 3-[(3-Cholamidopropyl)dimethylammonio]-1-propanesulfonate (CHAPS). This mixture was incubated at 4°C for ≥30 min. A 3.5 µL of sample was deposited onto 300 mesh Au-Flat 0.6/1.0 grids that had been glow discharged in a PELCO easiGLOW (Ted Pella) at 0.39 mBar, 30 mA for 60 s. Samples were vitrified in 100% liquid ethane using a Vitrobot Mark IV (Thermo Fisher Scientific), with a wait time of 2 s, blot time of 17 s, blot force of 5 at 100% humidity and 22°C. 350-341:H3 complex was prepared similarly to the S1V2-72:H3 complex, except the Fab:EHA2 complex was diluted to 4 mg/ml (final) before incubation at 4°C, and the grids were blotted for 11-20 s before vitrification.

For S1V2-72:B complex, B EHA2 was mixed with 1.5-fold molar excess of S1V2-72 Fab and diluted to 2 mg/ml protein in 10 mM Tris plus 150 mM NaCl (pH 8). This mixture was incubated on ice ≥30 min, then spiked with n-octyl-ß-D-glucoside (0.1% [w/v] final) so that the final protein concentration was 1.5 mg/ml. The final mixture was incubated at 4°C for 45 min. A 3.5 µL of sample was deposited onto 400 mesh HexAuFoil 0.3/0.3 grids that had been glow discharged at 0.39 mBar, 30 mA for 90 s. Samples were vitrified in 100% liquid ethane using a Vitrobot Mark IV, with a wait time of 2 s, blot time of 13 s, blot force of 5 at 100% humidity and 4°C.

### Cryo-EM data collection and processing

Cryo-EM data were recorded on a 300 kV Titan Krios G3i Microscope (Thermo Fisher Scientific) equipped with a Falcon 4i detector and a Selectris energy filter (10 eV slit width) at the Harvard Cryo-Electron Microscopy Center for Structural Biology at Harvard Medical School. Data were acquired in counting mode using the automated data collection software EPU (Thermo Fisher Scientific). Details of the data collection and dataset parameters are summarized in Table S2.

All data processing was performed using CryoSPARC (*55*). Dose-fractionated images were gain-normalized, aligned, dose-weighted and summed with patch motion correction, and then patch contrast transfer function (CTF) and defocus value estimation were performed.

The following particle-picking and processing strategy was used for each cryo-EM dataset. To generate initial templates for particle picking, the low-resolution atomic model of S1V2-72 Fab bound to B EHA2 (PDB: 8UDG) (*19*) was converted to a C3-symmetric model in UCSF ChimeraX (*56*). The ChimeraX molmap utility was used to construct a 16Å-resolution simulated volume from the C3-symmetric model. Fifty evenly spaced, projection-based templates were generated from this simulated volume in CryoSPARC. These templates were used for template-based particle picking in CryoSPARC. Picked particles were 2D-classified and the particles from the best-looking classes were used for ab-initio reconstruction of several classes. The 3D class with density clearly corresponding to three Fabs bound to EHA2 homotrimer was culled of junk particles by heterogenous refinement, and the refined volume was used to create new 2D templates for another round of particle picking. Junk particles were then culled by 2D classification and by heterogenous refinement of 3D volumes. For the final round of particle-picking, a selected set of particles were used to train a Topaz particle-picking model (*57*), which then picked new particles from all micrographs in the dataset. These particles were pooled with those retained after earlier rounds of picking and culling, then duplicate particles were removed, and additional rounds of heterogenous refinement were used to cull junk particles. The final particle set was subjected to non-uniform refinement (*58*) with reference-based motion correction (*59*) to produce the final EM map.

### Atomic model building and refinement

Structural biology applications used in this project were compiled and configured by SBGrid (*60*). ChimeraX was used to fit the atomic model of the complex of S1V2-72 Fab and B EHA2 (PDB: 8UDG) to the new, high-resolution density map for S1V2-72:B EHA2. The atomic model of S1V2-72 Fab (PDB: 8UDG) was also fit to the density map for S1V2-72:H3 EHA2. ModelAngelo (*61*) was used to build atomic models of the 350-341 Fab and H3 EHA2 into the EM density maps. Parts of the atomic models that were not built by ModelAngelo – but for which there was sufficient map density to justify placement of additional residues – were built manually in Coot (*62*) (v0.9.8.91). The constant domains of the Fabs were ignored, due to low density at these regions of the maps. The atomic models were refined with the ISOLDE plugin (*63*) of UCSF ChimeraX (*56*) (v1.9) and with PHENIX (*64, 65*) (v1.21.1-5286). Martin numbering was applied to the Fab residues with the Abnum server (*66*). PyMOL (*67*) (v3.1.8) and ChimeraX were used for visualization and analysis of the final model.

## STATISTICAL ANALYSES

GraphPad Prism (v10) software was used to perform statistical analyses as described in the figure legends.

## Author Contributions

J.F. and S.C.H. designed the research; J.F., J.P., M.K., D.T. performed the experiments; all authors analyzed the data.; J.F. and S.C.H. wrote the paper; all authors reviewed and approved the paper.

