## Supplemental_figures_and_tables for "Genetically restricted human antibodies recognize a conserved β-hairpin in postfusion influenza hemagglutinin"

**Figure S1. IgGs targeting the HA2 β-hairpin bind antigenically diverse IAV and IBV HAs.** A) Binding of recombinant IgGs to EHA2 representing IAV H1-H16 and IBV, or to trimeric, full-length soluble HA ectodomain (FLSE) representing H17 and H18, as measured by ELISA. B) IgG binding to N135R EHA2s, measured by ELISA.

**Figure S2. Single-particle cryo-EM analysis of the complex of S1V2-72 Fab and H3 EHA2.** A) Representative electron micrograph showing immunocomplex particles in vitreous ice. The image was low-pass filtered to 5 Å. Scale bar: 200 Å. B) 2-dimensional class averages of the particles included in the final cryo-EM map. C) Heatmap showing the distribution of viewing directions for particles included in the final cryo-EM map. D) Gold-standard Fourier Shell Correlation (GSFSC) curves of the final cryo-EM map. E) Final cryo-EM map, colored by local resolution. F) Cartoon model of the immunocomplex superposed with the cryo-EM map (transparent surface). G) Cartoon model of the homotrimeric immunocomplex. For clarity, only one protomer is colored.

**Figure S3. Single-particle cryo-EM analysis of the complex of S1V2-72 Fab and B EHA2.** A) Representative electron micrograph showing immunocomplex particles in vitreous ice. The image was low-pass filtered to 5 Å. Scale bar: 200 Å. B) 2-dimensional class averages of the particles included in the final cryo-EM map. C) Heatmap showing the distribution of viewing directions for particles included in the final cryo-EM map. D) Gold-standard Fourier Shell Correlation (GSFSC) curves of the final cryo-EM map. E) Final cryo-EM map, colored by local resolution. F) Cartoon model of the immunocomplex superposed with the cryo-EM map (transparent surface). G) Cartoon model of the homotrimeric immunocomplex. For clarity, only one protomer is colored.

**Figure S4. Structure of S1V2-72 Fab bound to postfusion B HA2.** A) Model of S1V2-72 Fab bound to B EHA2. Only one protomer of the Fab:EHA2 complex is colored. Subsequent panels indicate the rotations applied to the view in (A) to achieve the views shown in those panels. B) Interactions between the S1V2-72 HC and the HA2 β-hairpin. Black dashed lines, nonpolar interactions; magenta dashed lines, polar interactions. C) Interactions between the S1V2-72 LC and the HA2 β-hairpin, denoted as in (C). D) Interactions between the S1V2-72 HC and the ascending and descending α-helices of B EHA2. E) Sequence alignment of the S1V2-72 epitope residues (from the S1V2-72 Fab:B EHA2 complex) with the same residues from other IAV and IBV HA subtypes. F)   G) Left: Surface rendering of a model of prefusion HA (PDB: 6HJQ) next to a model of the S1V2-72 Fab:H3 EHA2 immunocomplex, where the EHA2 component is not displayed. One protomer of prefusion HA is colored teal, and the partly obscured HA2 β-hairpin of that protomer is colored red. Right: The HA and S1V2-72 models were aligned at the HA2 β-hairpin to dock the S1V2-72 Fab to the prefusion HA. A substantial steric clash between the S1V2-72 HC and an adjacent HA protomer is marked with a black oval.

**Figure S5. Single-particle cryo-EM analysis of the complex of 350-341 Fab and H3 EHA2.** A) Representative electron micrograph showing immunocomplex particles in vitreous ice. The image was low-pass-filtered to 5 Å. Scale bar: 200 Å. B) 2-dimensional class averages of the particles included in the final cryo-EM map. C) Heatmap showing the distribution of viewing directions for particles included in the final cryo-EM map. D) Gold-standard Fourier Shell Correlation (GSFSC) curves of the final cryo-EM map. E) Final cryo-EM map, colored by local resolution. F) Cartoon model of the immunocomplex superposed with the cryo-EM map (transparent surface). G) Cartoon model of the homotrimeric immunocomplex. For clarity, only one protomer is colored.

**Figure S6. Binding of β-hairpin Abs to H9, H10, and H11 is rescued by mutations that restore the consensus epitope.** A-C) Binding of IgGs to wildtype or mutant versions of H9 (A), H10 (B), or H11 (C) EHA2s, as measured by ELISA.

**Table S1. Features of candidate Abs that did not bind EHA2.**

| Antibody name | Donor | IGHV gene and allele | IGHD gene | IGHJ gene | HCDR3 | HCDR3 length | IGHV mutation frequency (%) | IGLV gene and allele | IGLJ gene and allele | LCDR3 | LCDR3 length | IGLV mutation frequency (%) | Reference |
|---|---|---|---|---|---|---|---|---|---|---|---|---|---|
| P05-7112 | P05 | HV1-2*04 | HD1-26 | HJ4 | ARASGGYYVLDN | 12 | 7.1 | LV2-11*01 | LJ3*02 | CSYAGSYTWV | 10 | 1.7 | McIntire et al. |
| P05-11077 | P05 | HV1-2*04 | HD3-16 | HJ4 | ARLAGRGSNAAY | 12 | 8.2 | LV2-11*01 | LJ3*02 | CSYAGTYTWV | 10 | 3.4 | McIntire et al. |
| P04-456 | P04 | HV1-2*02 | HD1-26 | HJ4 | ARIDIVGAHDY | 11 | 0.0 | LV2-14*01 | LJ3*02 | SSYTSSSTWV | 10 | 0.0 | McIntire et al. |
| P04-1084 | P04 | HV1-2*02 | HD2-8 | HJ6 | ARVVYFYYGVDV | 12 | 6.1 | LV2-14*01/*03 | LJ3*02 | NSYTSRSTWV | 10 | 3.7 | McIntire et al. |
| P04-1595 | P04 | HV1-2*02 | HD5-18 | HJ4 | ARDRRGLKHLWTY | 13 | 2.7 | LV2-14*01 | LJ3*02 | CSYTTNSTWV | 10 | 2.7 | McIntire et al. |
| P04-2825 | P04 | HV1-2*02 | HD1-26 | HJ4 | ARDPGGRYEIDY | 12 | 1.7 | LV2-14*01 | LJ3*02 | SSYTSSSTWV | 10 | 1.0 | McIntire et al. |
| P04-2866 | P04 | HV1-2*02 | HD1-1 | HJ4 | ASQSASRPELDY | 12 | 4.8 | LV2-14*04 | LJ3*02 | SSYTSSSTWV | 10 | 1.0 | McIntire et al. |
| P04-3803 | P04 | HV1-2*02 | HD5-18 | HJ4 | ATGSTYGWGAEGY | 13 | 8.2 | LV2-14*01 | LJ3*02 | ISYTSTGTSWV | 11 | 6.1 | McIntire et al. |
| P04-4501 | P04 | HV1-2*02 | HD6-6 | HJ3 | ARGRYTSSPFKV | 12 | 4.4 | LV2-14*01/*03 | LJ3*02 | SSFTSSTSWV | 10 | 4.8 | McIntire et al. |
| P05-6366 | P05 | HV1-2*04 | HD2-8 | HJ4 | ARFYFRSLDY | 10 | 4.8 | LV2-14*03/*04 | LJ3*02 | GSYTGNTVWV | 10 | 6.6 | McIntire et al. |
| P05-6540 | P05 | HV1-2*04 | HD1-26 | HJ4 | ARGGSNYRSPFDY | 13 | 2.0 | LV2-14*04 | LJ3*02 | SSYTSNNRWV | 10 | 2.4 | McIntire et al. |
| P05-6638 | P05 | HV1-2*04 | HD6-13 | HJ5 | AREVTSSWFDY | 11 | 3.0 | LV2-14*04 | LJ3*02 | ISYTSSSTWV | 10 | 1.7 | McIntire et al. |
| P05-7024 | P05 | HV1-2*04 | HD5-18 | HJ3 | ARDGCRCHAFDV | 12 | 7.1 | LV2-14*03 | LJ3*02 | SSYTNTNTWV | 10 | 6.5 | McIntire et al. |
| P05-7078 | P05 | HV1-2*04 | HD2-15 | HJ4 | ARGSSWNYFDY | 11 | 0.3 | LV2-14*03 | LJ3*02 | SSYTSSTTWV | 10 | 0.3 | McIntire et al. |
| P05-7355 | P05 | HV1-2*04 | HD6-19 | HJ4 | ARVLRSSGRLYP | 12 | 0.0 | LV2-14*04 | LJ3*02 | SSYTSSSHWV | 10 | 0.3 | McIntire et al. |
| P05-7451 | P05 | HV1-2*04 | HD4-23 | HJ3 | ARNYGDIRRGAFDI | 14 | 0.3 | LV2-14*04 | LJ3*02 | SSYTNSSTWV | 10 | 1.4 | McIntire et al. |
| P05-7567 | P05 | HV1-2*04 | HD3-10 | HJ2 | ARAPRGSGRHFDL | 13 | 0.0 | LV2-14*04 | LJ3*02 | SSYTSSSTWV | 10 | 0.3 | McIntire et al. |
| P11-8541 | P11 | HV1-2*04 | HD1-14 | HJ6 | VRDSGNYGMDV | 11 | 9.1 | LV2-14*03 | LJ3*02 | SSYTRSTSWV | 10 | 5.8 | McIntire et al. |
| P11-8597 | P11 | HV1-2*04 | HD7-27 | HJ3 | AAELDNWGSNAFDV | 14 | 4.5 | LV2-14*03 | LJ3*02 | SSYTTSNSWV | 10 | 3.1 | McIntire et al. |
| P05-6545 | P05 | HV1-2*04 | HD3-16 | HJ1 | AREYGRGAYPD | 11 | 1.0 | LV2-23*04 | LJ3*02 | CSYAGSSTWV | 10 | 1.4 | McIntire et al. |
| C4-21455 | C4 | HV1-2*06 | HD1-14 | HJ4 | ARAGGEPTAPDY | 12 | 3.1 | LV2-23*02 | LJ3*02 | CSYTGSSTFKV | 11 | 3.0 | Park et al. |

**Table S2: Cryo-EM data collection, refinement and validation statistics**

| | **S1V2-72 : B EHA2**<br>PDB: 12RO<br>EMDB: EMD-76712 | **S1V2-72 : H3 EHA2**<br>PDB: 12RK<br>EMDB: EMD-76710 | **350-341 : H3 EHA2**<br>PDB: 12RB<br>EMDB: EMD-76704 |
|---|---|---|---|
| **Data collection and processing** | | | |
| Magnification | 215,000× | 165,000× | 165,000× |
| Voltage (kV) | 300 | 300 | 300 |
| Electron exposure ($e^{-}$/Å$^{-2}$) | 50.49 | 53.57 | 52.82 |
| Defocus range (μm) | -0.8, -2.0 | -0.8, -2.0 | -0.8, -2.0 |
| Pixel size (Å) | 0.579 | 0.737 | 0.736 |
| Symmetry imposed | C3 | C3 | C3 |
| Final particle images (no.) | 181,041 | 101,878 | 48,366 |
| Map Resolution (Å) | 2.95 | 3.14 | 3.47 |
| FSC threshold | 0.143 | 0.143 | 0.143 |
| Map resolution range (Å) | 2.5 - 8.0 | 2.8 - 10.0 | 3.0 - 12 |
| **Refinement** | | | |
| Initial model used | 8UDG | Fab: 8UDG<br>EHA2: de novo<br>(ModelAngelo) | Fab & EHA2: de novo<br>(ModelAngelo) |
| Map-sharpening B-factor (Å$^2$) | -79.8 | -83.5 | -91.7 |
| Model composition | | | |
| Non-hydrogen atoms | 8,397 | 8,667 | 8,718 |
| Protein residues | 1,101 | 1,104 | 1,107 |
| Waters | 0 | 6 | 3 |
| B factors (Å$^2$) | | | |
| Protein (min/max/mean) | 1.20/124.08/52.49 | 7.56/115.98/61.46 | 20.99/101.87/54.91 |
| R.M.S.D. deviations | | | |
| Bond lengths (Å) (# > 4σ) | 0.005 (0) | 0.007 (0) | 0.005 (0) |
| Bond angles (°) (# > 4σ) | 1.106 (0) | 1.104 (0) | 1.001 (0) |
| **Validation** | | | |
| MolProbity score | 1.36 | 0.97 | 1.21 |
| Clashscore | 4.35 | 2.00 | 3.16 |
| EM-Ringer score | 3.25 | 4.36 | 4.01 |
| Model-map correlation coefficient (mask) | 0.72 | 0.80 | 0.80 |
| rotamer outliers (%) | 0.32 | 0.63 | 0.32 |
| Ramachandran plot | | | |
| Favored (%) | 97.23 | 98.07 | 97.52 |
| Allowed (%) | 2.77 | 1.93 | 2.48 |
| Disallowed (%) | 0.00 | 0.00 | 0.00 |

**Fig. S1**

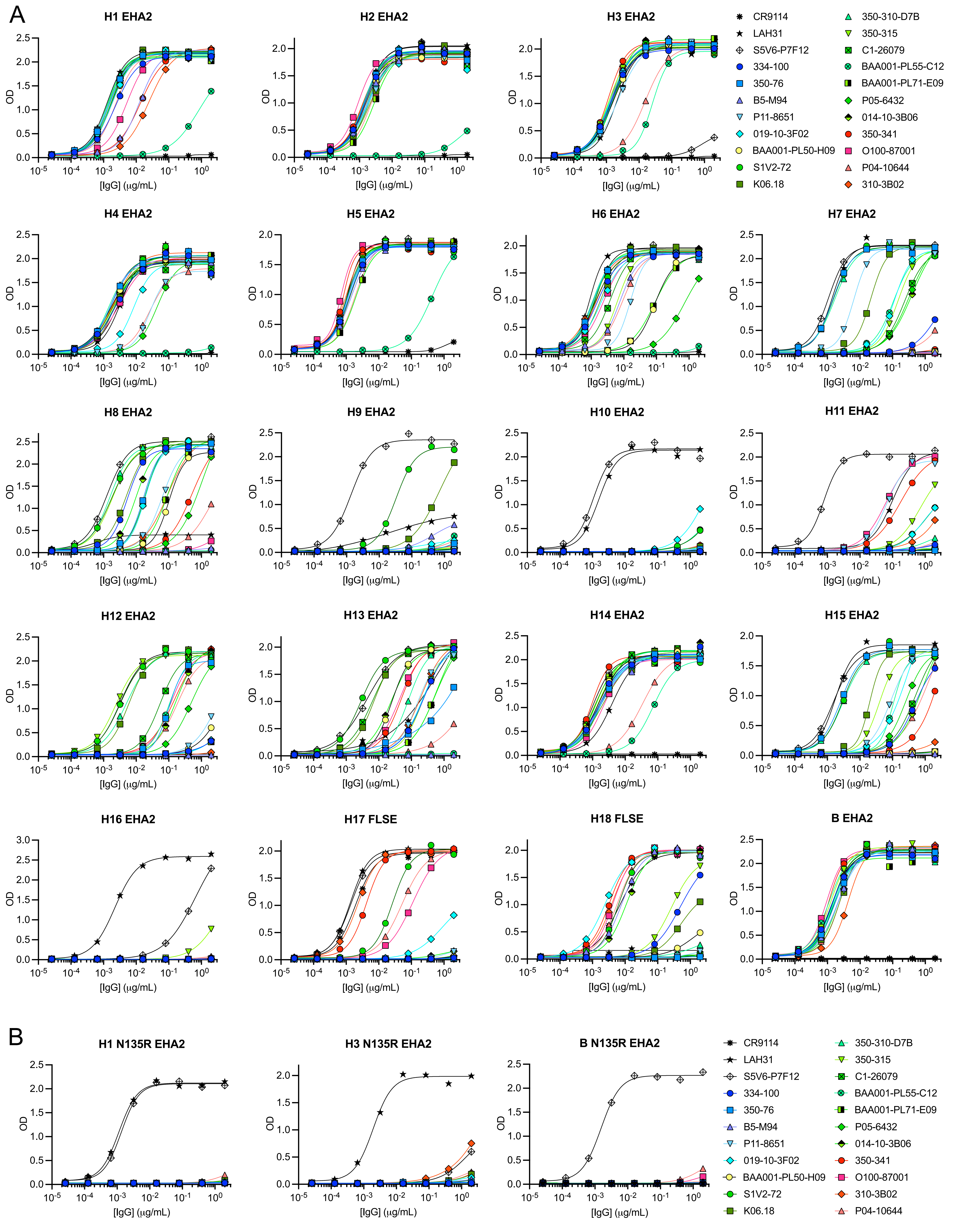
A
H1 EHA2
H2 EHA2
H3 EHA2
H4 EHA2
H5 EHA2
H6 EHA2
H7 EHA2
H8 EHA2
H9 EHA2
H10 EHA2
H11 EHA2
H12 EHA2
H13 EHA2
H14 EHA2
H15 EHA2
H16 EHA2
H17 FLSE
H18 FLSE
B EHA2
OD
[IgG] (μg/mL)
CR9114
LAH31
S5V6-P7F12
334-100
350-76
B5-M94
P11-8651
019-10-3F02
BAA001-PL50-H09
S1V2-72
K06.18
350-310-D7B
350-315
C1-26079
BAA001-PL55-C12
BAA001-PL71-E09
P05-6432
014-10-3B06
350-341
O100-87001
P04-10644
310-3B02
B
H1 N135R EHA2
H3 N135R EHA2
B N135R EHA2

Fig. S2

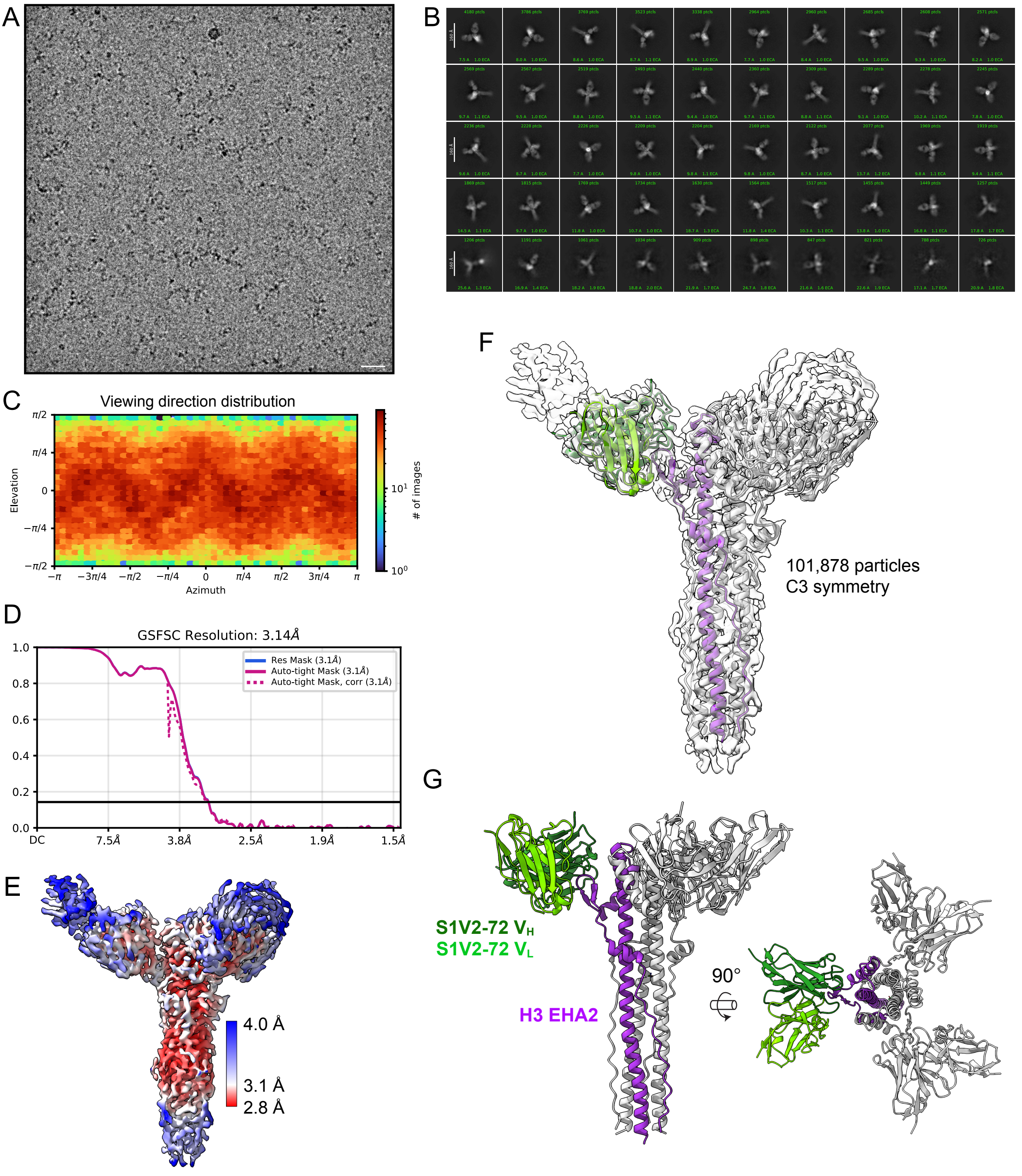
A
B
C
Viewing direction distribution
Elevation
Azimuth
### of images
D
GSFSC Resolution: 3.14Å
Res Mask (3.1Å)
Auto-tight Mask (3.1Å)
Auto-tight Mask, corr (3.1Å)
DC
7.5Å
3.8Å
2.5Å
1.9Å
1.5Å
E
4.0 Å
3.1 Å
2.8 Å
F
101,878 particles
C3 symmetry
G
S1V2-72 $V_H$
S1V2-72 $V_L$
H3 EHA2
90°

Fig. S3

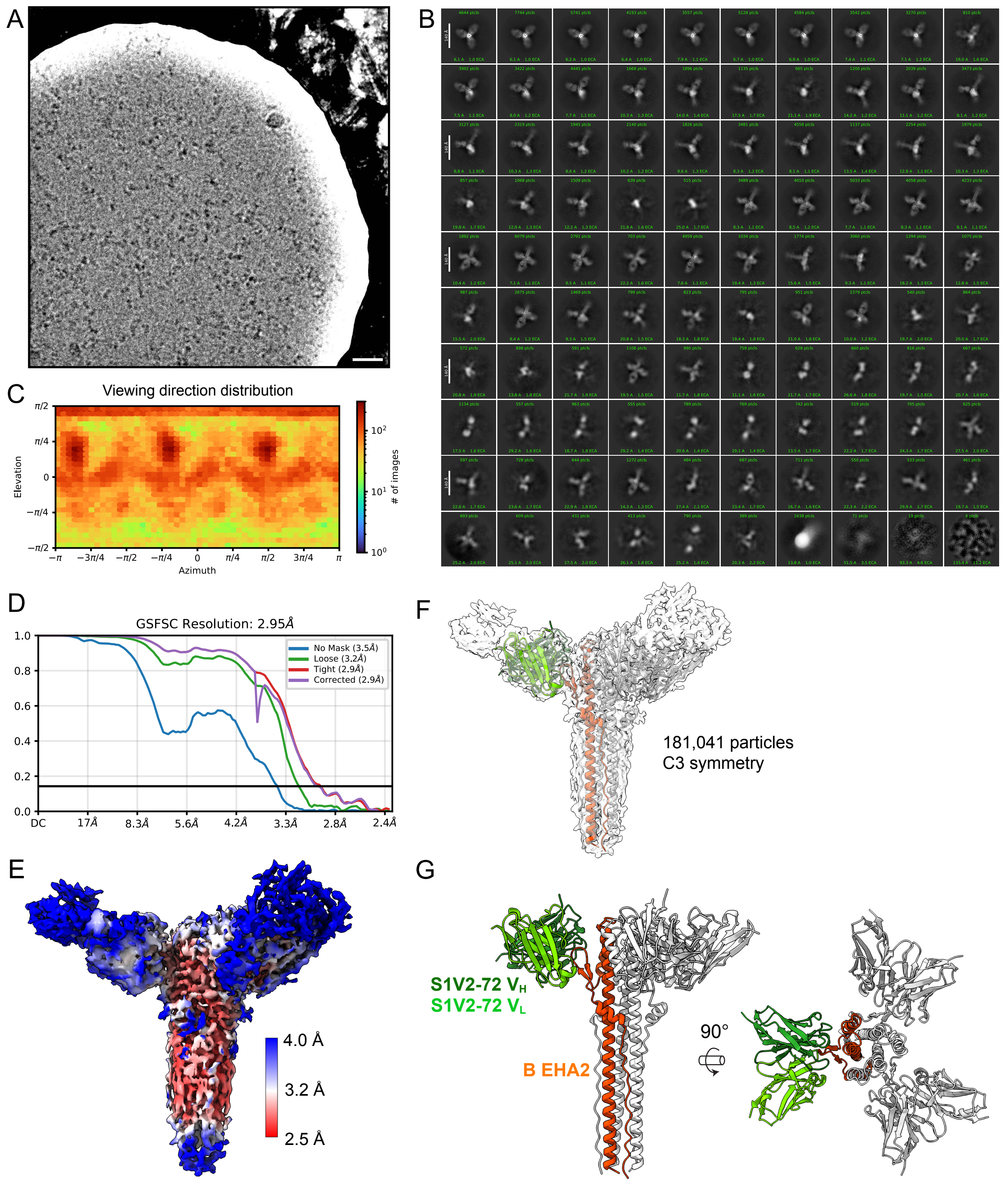

A
B
C
Viewing direction distribution
Elevation
Azimuth
### of images
D
GSFSC Resolution: 2.95Å
No Mask (3.5Å)
Loose (3.2Å)
Tight (2.9Å)
Corrected (2.9Å)
DC
17Å
8.3Å
5.6Å
4.2Å
3.3Å
2.8Å
2.4Å
E
4.0 Å
3.2 Å
2.5 Å
F
181,041 particles
C3 symmetry
G
S1V2-72 V$_H$
S1V2-72 V$_L$
B EHA2
90°

Fig S4

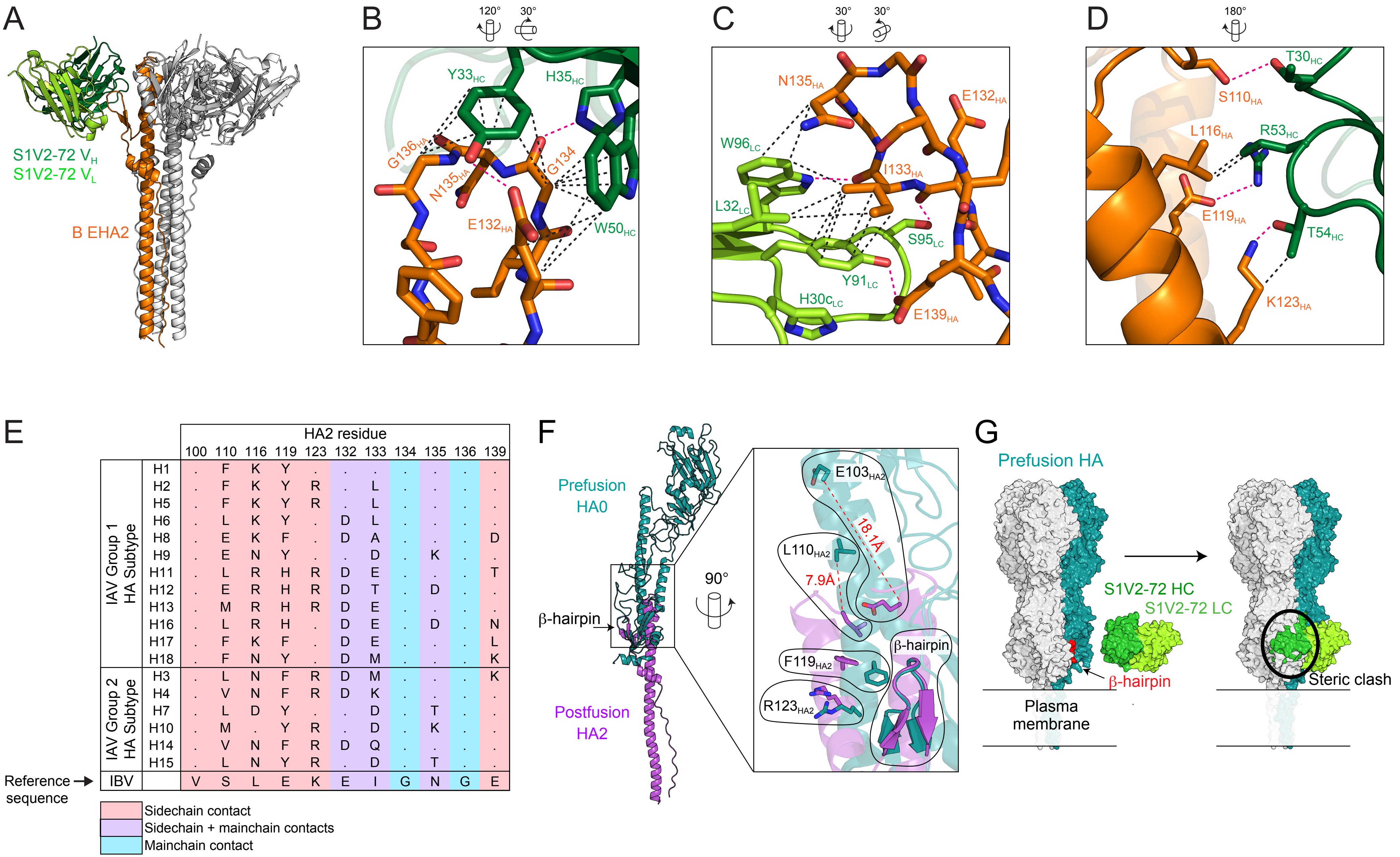


| | | HA2 residue 100 | 110 | 116 | 119 | 123 | 132 | 133 | 134 | 135 | 136 | 139 |
|---|---|---|---|---|---|---|---|---|---|---|---|---|
| IAV Group 1 HA Subtype | H1 | . | F | K | Y | . | . | . | . | . | . | . |
| | H2 | . | F | K | Y | R | . | L | . | . | . | . |
| | H5 | . | F | K | Y | R | . | L | . | . | . | . |
| | H6 | . | L | K | Y | . | D | L | . | . | . | . |
| | H8 | . | E | K | F | . | D | A | . | . | . | D |
| | H9 | . | E | N | Y | . | . | D | . | K | . | . |
| | H11 | . | L | R | H | R | D | E | . | . | . | T |
| | H12 | . | E | R | H | R | D | T | . | D | . | . |
| | H13 | . | M | R | H | R | D | E | . | . | . | . |
| | H16 | . | L | R | H | . | D | E | . | D | . | N |
| | H17 | . | F | K | F | . | D | E | . | . | . | L |
| | H18 | . | F | N | Y | . | D | M | . | . | . | K |
| IAV Group 2 HA Subtype | H3 | . | L | N | F | R | D | M | . | . | . | K |
| | H4 | . | V | N | F | R | D | K | . | . | . | . |
| | H7 | . | L | D | Y | . | . | D | . | T | . | . |
| | H10 | . | M | . | Y | R | . | D | . | K | . | . |
| | H14 | . | V | N | F | R | D | Q | . | . | . | . |
| | H15 | . | L | N | Y | R | . | D | . | T | . | . |
| IBV | | V | S | L | E | K | E | I | G | N | G | E |

Fig. S5

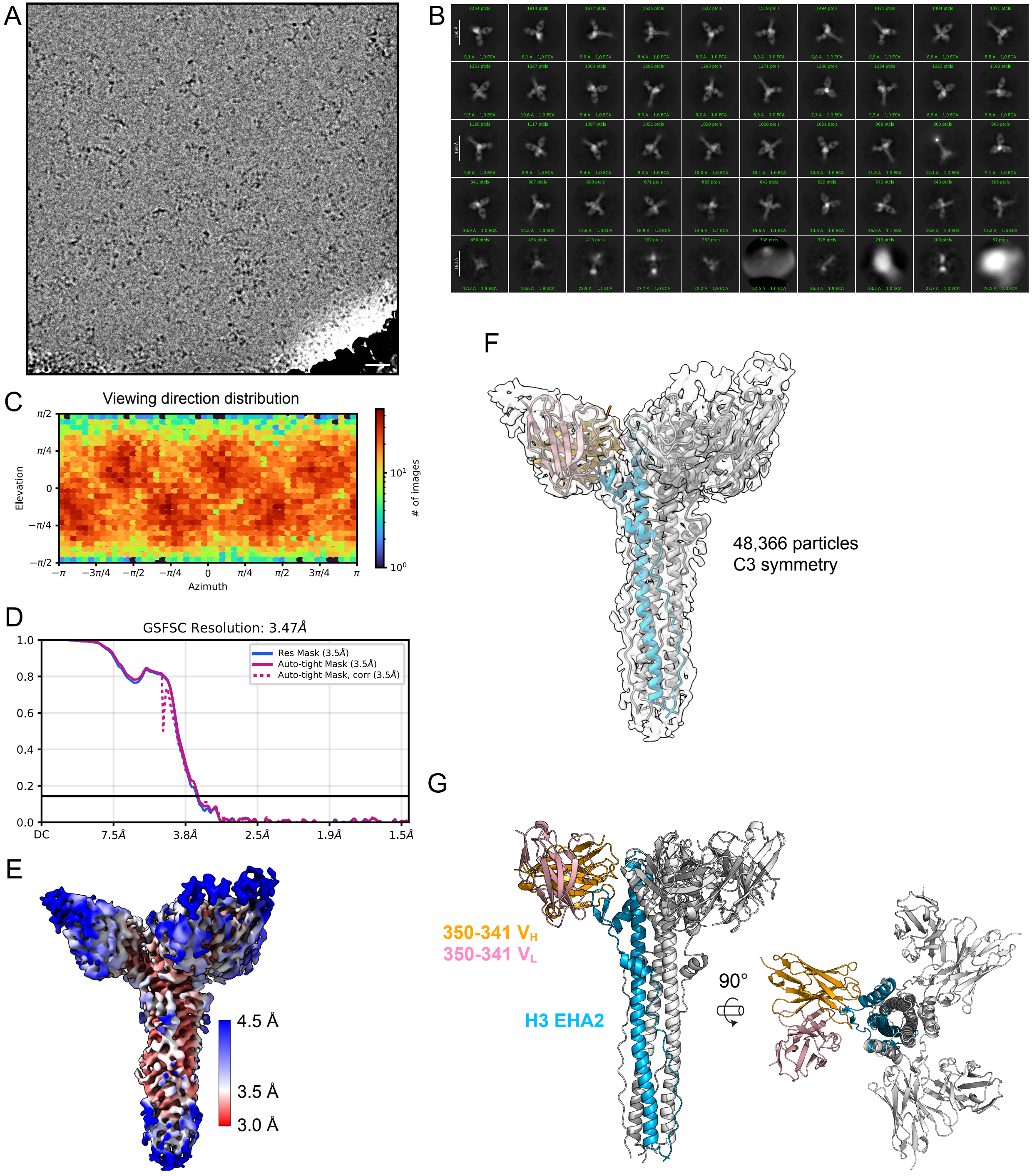
A
B
C
Viewing direction distribution
Elevation
Azimuth
### of images
D
GSFSC Resolution: 3.47Å
Res Mask (3.5Å)
Auto-tight Mask (3.5Å)
Auto-tight Mask, corr (3.5Å)
DC
7.5Å
3.8Å
2.5Å
1.9Å
1.5Å
E
4.5 Å
3.5 Å
3.0 Å
F
48,366 particles
C3 symmetry
G
350-341 V_H
350-341 V_L
H3 EHA2
90°

**Fig. S6**

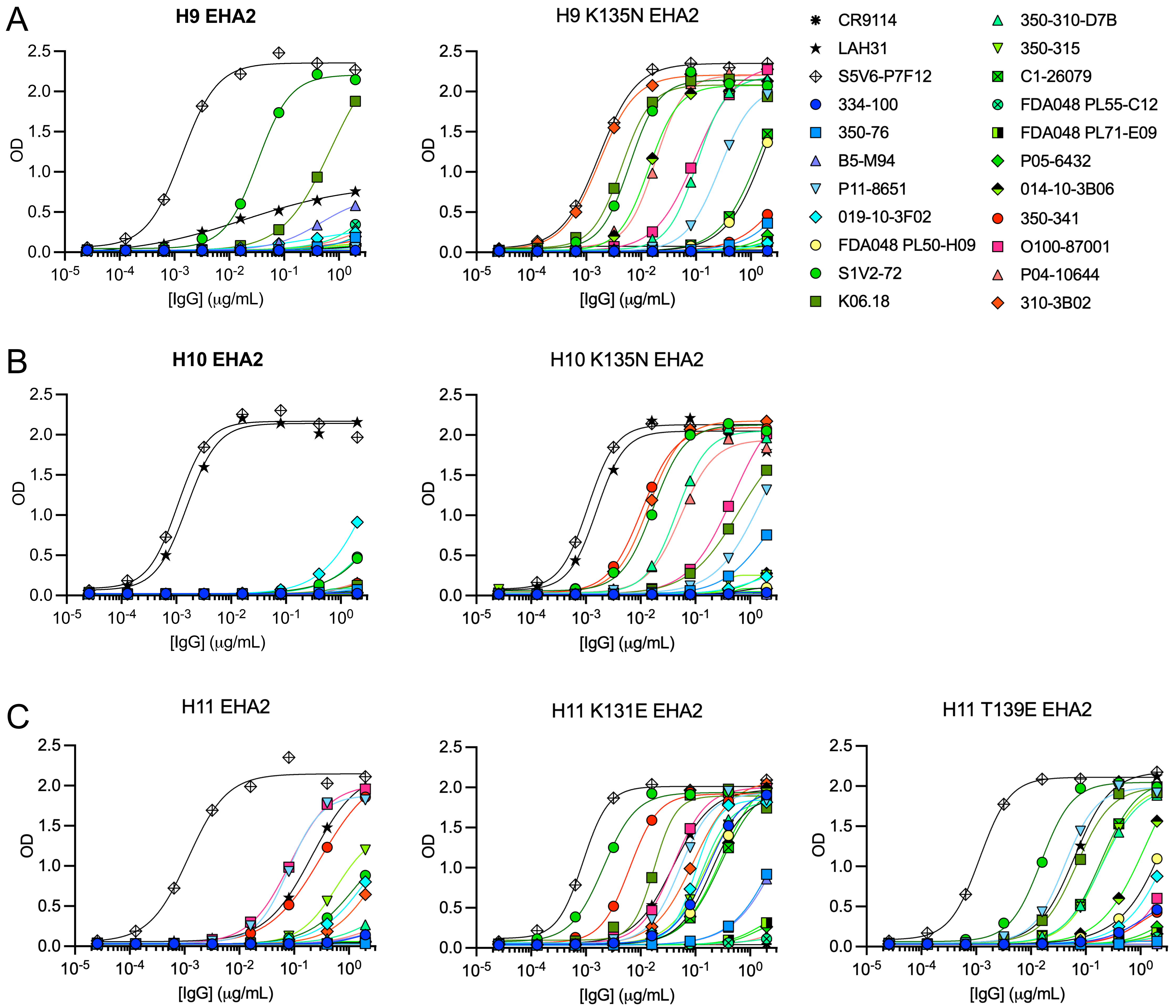
A
H9 EHA2
H9 K135N EHA2
B
H10 EHA2
H10 K135N EHA2
C
H11 EHA2
H11 K131E EHA2
H11 T139E EHA2
OD
[IgG] (μg/mL)
CR9114
LAH31
S5V6-P7F12
334-100
350-76
B5-M94
P11-8651
019-10-3F02
FDA048 PL50-H09
S1V2-72
K06.18
350-310-D7B
350-315
C1-26079
FDA048 PL55-C12
FDA048 PL71-E09
P05-6432
014-10-3B06
350-341
O100-87001
P04-10644
310-3B02
